# Purinergic receptor activation rectifies autism-associated endothelial dysfunction

**DOI:** 10.1101/2025.06.02.657522

**Authors:** Julie Ouellette, Sareen Warsi, Phinea Romero, Purva Khare, Shama Naz, Leya Aubert-Tandon, Chantal Pileggi, Sozerko Yandiev, Moises Freitas-Andrade, Cesar H. Comin, Mary-Ellen Harper, Devika S Manickam, Fabrice Dabertrand, Armen Saghatelyan, Baptiste Lacoste

**Affiliations:** Neuroscience Program, The Ottawa Hospital Research Institute, Ottawa, ON, Canada; Cellular & Molecular Medicine, University of Ottawa, Ottawa, ON, Canada; Departments. of Anesthesiology and of Pharmacology, University of Colorado Anschutz Medical Campus, CO, USA; Graduate School of Pharmaceutical Sciences, Duquesne University, Pittsburgh, PA, USA; University of Ottawa Metabolomics Core Facility, Faculty of Medicine, Ottawa, ON, Canada; Department of Biochemistry Microbiology and Immunology, Faculty of Medicine, University of Ottawa, Ottawa, Ontario, Canada; Federal University of São Carlos, Department of Computer Science, São Carlos, SP, Brazil; University of Ottawa Brain and Mind Research Institute, Ottawa, ON, Canada

**Author notes:** Correspondence to: Dr. Baptiste Lacoste, Faculty of Medicine, Department of Cellular and Molecular Medicine University of Ottawa & Ottawa Hospital Research Institute, 451 Smyth Road, Ottawa (ON) K1H 8M5, Ottawa, Canada.

**Keywords:** Endothelium, autism, metabolism, ATP, postnatal, brain, angiogenesis, mice, P2 receptors

## Abstract

Early cerebrovascular alterations affect brain maturation by impacting trophic support and energy supply. Recent evidence in a 16p11.2 deletion mouse model of autism spectrum disorder (ASD) revealed brain endothelial abnormalities postnatally. Yet, the endothelial alterations eliciting these changes remain unknown. Isolation of brain endothelial cells (ECs) from 14-day old male 16p11.2-deficient and wild-type mice revealed that 16p11.2 deletion-induced endothelial dysfunction is linked to a bioenergetic failure, with reduced intracellular ATP. Intra- or extra-cellular ATP supplementation rescued the function of 16p11.2-deficient ECs *in vitro* via P2 purinergic receptor activation, specifically P2Y2 receptors. Activating P2Y2 receptors restored cerebrovascular reactivity in 16p11.2-deficient parenchymal arterioles *ex vivo* and rescued 16p11.2 deletion-associated mouse behaviors. Taken together, this study demonstrates that metabolic reprogramming of brain ECs via purinergic receptor engagement represents a possible therapeutic avenue for ASD.

## Main

The brain is highly reliant on metabolic health, which includes uninterrupted delivery of nutrients and oxygen from its vasculature^1^. Energy requirements during critical periods of development further increase the vulnerability of the brain to metabolic failures^2,3^. During development, brain endothelial cells (ECs) help guide and coordinate neurogenesis, while neuronal cues regulate vascularization of the central nervous system^4^. As such, brain endothelial dysfunction can compromise metabolic support to neuronal cells, leading to the progression and/or onset of neurological conditions. Recent evidence revealed a causal link between altered brain EC function and autism spectrum disorders (ASD) in a mouse model of the 16p11.2 deletion syndrome, a mutation commonly found in human ASD^5–8^. While ASD are generally depicted by neuronal features that are highly contingent on brain metabolism^9–15^, the recent association with endothelial dysfunction warrants further inquiry. Here, we investigated the impact of a 16p11.2 deletion on brain EC metabolism at a key milestone of cerebrovascular development (post-natal day 14) to unveil new mechanisms underlying ASD pathophysiology. We demonstrate that 16p11.2 deletion-induced endothelial dysfunction is caused by a bioenergetic failure, with reduced intracellular ATP. Activation of ATP signaling via endothelial P2-class purinergic receptors, more specifically the P2Y2 receptor, rescued EC dysfunction by restoring angiogenic capacity *in vitro* and endothelium-dependent cerebrovascular reactivity *ex vivo* in 16p11.2-deficient cortical parenchymal arterioles. Finally, a selective pharmacological P2Y2 agonist rescued adult 16p11.2-deficient behavioral phenotypes. Altogether, these findings open new avenues for the development of therapies for ASD.

### Bioenergetic failure in *16p11.2^df/+^* brain endothelial cells

Emergence of ASD-related phenotypes coinciding with endothelial dysfunction in 16p11.2 deletion^6^ has prompted us to focus on 16p11.2-deficient ECs *in vitro* parameters as a screening platform. We first confirmed that 16p11.2-deficient ECs isolated from 14-day old mice, a landmark of cerebrovascular maturation^16^, do not establish a vascular network *in vitro* as evidenced by absence of endothelial tubes following seeding of 16p11.2-deficient ECs in an extracellular matrix (Figure S1A). Our prior work also identified reduced mitochondria number in adult 16p11.2-deficient brain ECs *in vivo*^7^. We thus tested whether mitochondrial density was reduced *in vitro* in P14 brain ECs isolated from *16p11.2^df/+^* male mice and found a similar phenotype, without apparent change in mitochondrial fragmentation (Figure S1B). This suggests that mitochondrial discrepancies are present before adulthood and may impact EC function early.

Despite reduced mitochondrial network density in *16p11.2^df/+^*ECs at P14, levels of proteins involved in mitochondrial fusion (MFN1, MFN2 and OPA1) and fission (DRP1), as well as mitochondrial biogenesis (PGC-1α), appeared unchanged (Figure S1C). These findings lead us to hypothesize that 16p11.2 haploinsufficiency may cause metabolic alterations in P14 mouse brain ECs.

To test this, we investigated the bioavailability of intracellular metabolites in P14 brain ECs from *16p11.2^df/+^* and WT male mice by performing targeted metabolomics analysis. Distinct metabolite profiles were identified between *16p11.2^df/+^* and WT brain ECs, as shown by cluster separation following a partial least square-discriminant analysis (PLS-DA) (Figure S1D). We measured overall decreased metabolite abundance in *16p11.2^df/+^* ECs compared to WT ECs, which was consistent in two independent cohorts (Figure 1A). With no exception, changes observed were reductions in metabolite abundance in *16p11.2^df/+^* ECs (Figure 1B and Figure S1E). Remarkably, the top 20 metabolites driving cluster separation comprised high energy molecules including ATP, as well as co-factors involved in glycolysis and energy utilization (e.g., glutamine, pyruvic acid and aspartic acid) (Figure S1G). These results indicate that energy metabolism is altered in 16p11.2-deficient ECs. Specifically, a bioenergetic failure in 16p11.2-deficient ECs was evidenced by 50% reduction in abundance of ATP and by lower abundance of metabolites including high energy donors (UDP-glucose/UDP galactose), co-enzymes required for the Krebs cycle and oxidative phosphorylation (NAD^+^, FAD), and amino acids essential for balancing energy dynamics (alanine/sarcosine and creatine) (Figures S1F,H and S2A). *16p11.2^df/+^* ECs also displayed a significant reduction in both ATP:ADP ratio and adenylate energy charge, indicators of cellular energy status (Figure 1C). Notably, the reduced ATP in 16p11.2-deficient ECs was not attributed to changes in CD39 protein level, a rate-limiting enzyme involved in the breakdown cascade of ATP (Figures 1D and S1J). Moreover, no difference was found in the reduced glutathione-to-oxidized glutathione ratio (GSH:GSSG) between *16p11.2^df/+^* and WT ECs (Figure S1H), indicating that changes in metabolite abundance were not attributed to poor cellular health at the time of the metabolite extraction^17,18^, which is consistent with our previous work showing normal baseline endothelial health (*i.e.*, survival, proliferation and migration) in P14 *16p11.2^df/+^*brain ECs^6^. Although no difference was seen in the GSH:GSSG ratio, both GSH and GSSG were individually reduced in *16p11.2^df/+^* ECs compared to WT ECs, consistent with decreased gamma-glu-cys, a metabolite upstream of the glutathione pathway^19^ (Figures S1F and H). Unexpectedly, no difference was measured in ATP-sensing AMP-activated protein kinase (AMPK)^20^ activity between 16p11.2-deficient and WT ECs, despite 50% reduction in ATP abundance and a lower ATP:ADP ratio in 16p11.2-deficient ECs (Figure 1C and Figure S1I). Of note, abundance of the main regulators of AMPK activity (*i.e.,* AMP and ADP) were not significantly changed in 16p11.2-deficient ECs (Figure S1H).

**Figure 1.**
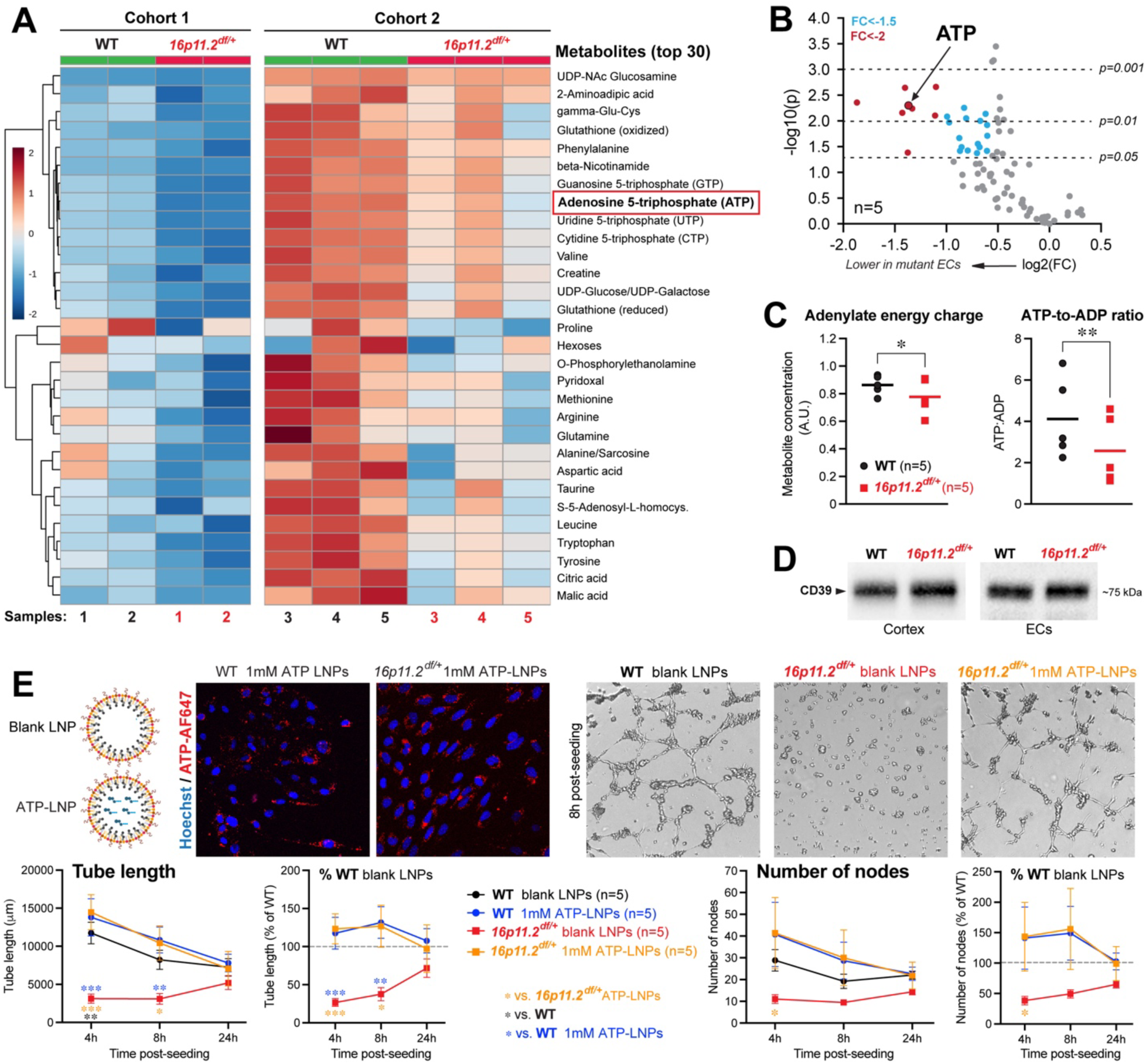
Primary 16p11.2-deficient brain endothelial cells isolated from P14 mice exhibit a bioenergetic failure and intracellular delivery of ATP rescues their angiogenic capacity. (A) Heatmap (multivariate statistical analysis) of two distinct cohorts displaying the top 30 metabolites significantly altered by genotype. (B) Volcano plot showing most significantly altered metabolites (*p*<0.01), including ATP, with a fold change (FC)<-2 (red) and FC<-1.5 (blue). Plot summarizes both fold-change and t-test criteria. Scatter-plot of the negative log10-transformed p-values from the t-test plotted against the log2 fold change is shown. (C) Quantitative assessment of energy-related metabolite ratio (adenylate energy charge and ATP-to-ADP ratio, n=5 samples). Data represents mean individual values. \**p*<0.05, \*\**p*<0.01, a two tailed paired *t-test*. (D) Representative Western blots of relative CD39 protein levels from 16p11.2-deficient and WT primary brain ECs or cerebral cortex (see Figure S1J for details). (E) *In vitro* network formation assay was used to assess angiogenic activity over 24 hours in a growth-factor-reduced Matrigel^®^. Top left: schematic representation of blank LNPs (without ATP) and ATP-LNP structures. Top middle: Representative images confirming intracellular delivery of ATP (red) via lipid nanoparticles (LNPs). Top right: representative images at the 8h post-seeding time point of WT and 16p11.2-deficient ECs treated with blank LNPs and 16p11.2-deficient ECs treated with 1mM of ATP-LNPs during a network formation assay. Bottom: quantifications of network densities (total endothelial tube length) and network nodes (total number of branching hubs) with representative percent of untreated WT. Dotted line represents 100% of untreated WT values. Data represents mean ± s.e.m. (n=5 animals per group). \**p*<0.05, \*\**p*<0.01, \*\**p*<0.001, two-way repeated measure ANOVA and Sidak’s multicomparison *post hoc* test. (A-E) All data shown are from ECs isolated from P14 male mice. (A-C) Data are from ECs pooled from 3 mice per sample (n=5 samples).

To test whether the observed bioenergetic failure was related to alterations in mitochondrial function, we used a Seahorse Cell Mito stress test. No difference was measured between P14 WT and *16p11.2^df/+^*ECs in oxygen consumption rate (OCR), extracellular acidification rate (ECAR), as well as additional parameters of mitochondrial function including ATP production (Figure S2B). These findings suggest that the bioenergetic failure evidenced in 16p11.2-deficient ECs might be a direct contributor to endothelial dysfunction, independent of the function of individual mitochondria. While at this stage the direct cause of reduced intracellular ATP in 16p11.2-deficient ECs remain to be elucidated, it raises the possibility of restoring ATP bioavailability as a rescue strategy.

### Intracellular delivery of ATP rescues *16p11.2^df/+^* endothelial dysfunction

In light of recent work showing that mitochondrial ATP is required to maintain cerebrovascular tone^21^, we sought to test whether intracellular ATP supplementation could normalize the cell-autonomous angiogenic capacity of 16p11.2-deficient ECs in a Matrigel®-based assay^6^. Here, we opted to deliver ATP intracellularly in cultured ECs via lipid nanoparticles (LNPs)^22^. Since physico-chemical properties of nanoparticles, such as colloidal stability, determine their biological efficiency^23,24^, we confirmed colloidally stable dispersions (particle sizes <200 nm and dispersity indices ∼0.3) of ATP-loaded LNPs (ATP-LNPs) over 14 days upon intermittent storage at 2-8°C as well as their sphere-like morphology (Figures S3A-C). The cellular uptake of LNPs containing AF647-labeled ATP^22^ compared to blank LNPs was also confirmed using fluorescence microscopy (Figure 1E and Figure S3D-E). 16p11.2-deficient and WT ECs were then treated with 1mM of ATP-LNPs from cell isolation for 7 days (6 days in culture and 24-hour tube formation assay) (Figure S3F). 16p11.2-deficient ECs treated with 1mM of ATP-LNPs displayed a dramatic increase in tube length (*i.e.,* branches) at the 4- and 8-hour timepoints compared to *16p11.2^df/+^* ECs treated with blank-LNPs (Figure 1E). These results demonstrate that intracellular delivery of ATP can boost the angiogenic capacity of 16p11.2-deficient brain ECs *in vitro*. Interestingly, increasing ATP-LNP concentration to 5mM did not confer rescue, but hindered the ability of *16p11.2^df/+^* ECs to establish a network (Figure S3G). These data support that 16p11.2 deletion-induced endothelial dysfunction can be rescued by intracellular ATP supplementation below a concentration threshold.

### P2-class purinergic receptor activation rescues 16p11.2 deletion endothelial dysfunction

The question remains what mechanisms underlie ATP-mediated rescue. ATP is known to act as a signaling molecule on brain ECs^25^, which are sensitive to ATP via expression of P2-class purinergic receptors^26^. Recent work demonstrated that intracellular ATP can exit brain ECs through pannexin-1 (Panx1) channels to subsequently activate membrane P2 purinergic receptors on adjacent ECs to regulate vascular tone^27^ (Figure 2A). We thus tested whether ATP delivered intracellularly via LNPs exit ECs via Panx1 channels to subsequently act on membrane P2 receptors. Notably, P14 mouse brain ECs expressed *Panx1* and produced Panx1 (Figure 2B). We also find that expression of *Panx1* declined towards adulthood (Figure 2B), consistent with previous work^28^, suggesting a role for Panx1 during postnatal development. Treatment of 16p11.2-deficient ECs with 1mM of a Panx1 antagonist, probenecid, prevented the 1mM ATP-LNP-mediated rescue of 16p11.2-deficient brain EC function as shown by a lack of tube formation in 1mM probenecid and 1mM ATP-LNP treated 16p11.2-deficient ECs compared to 1mM ATP-LNP treated 16p11.2-deficient ECs (Figure 2C). Of note, we found that probenecid treatment of 16p11.2-deficient ECs at 150μM or 500μM was also able to prevent network formation, however the most efficient dose of probenecid being 1mM (Figure S6A-D). This suggests that ATP exit ECs via Panx1 channels to exhibit its function.

**Figure 2.**
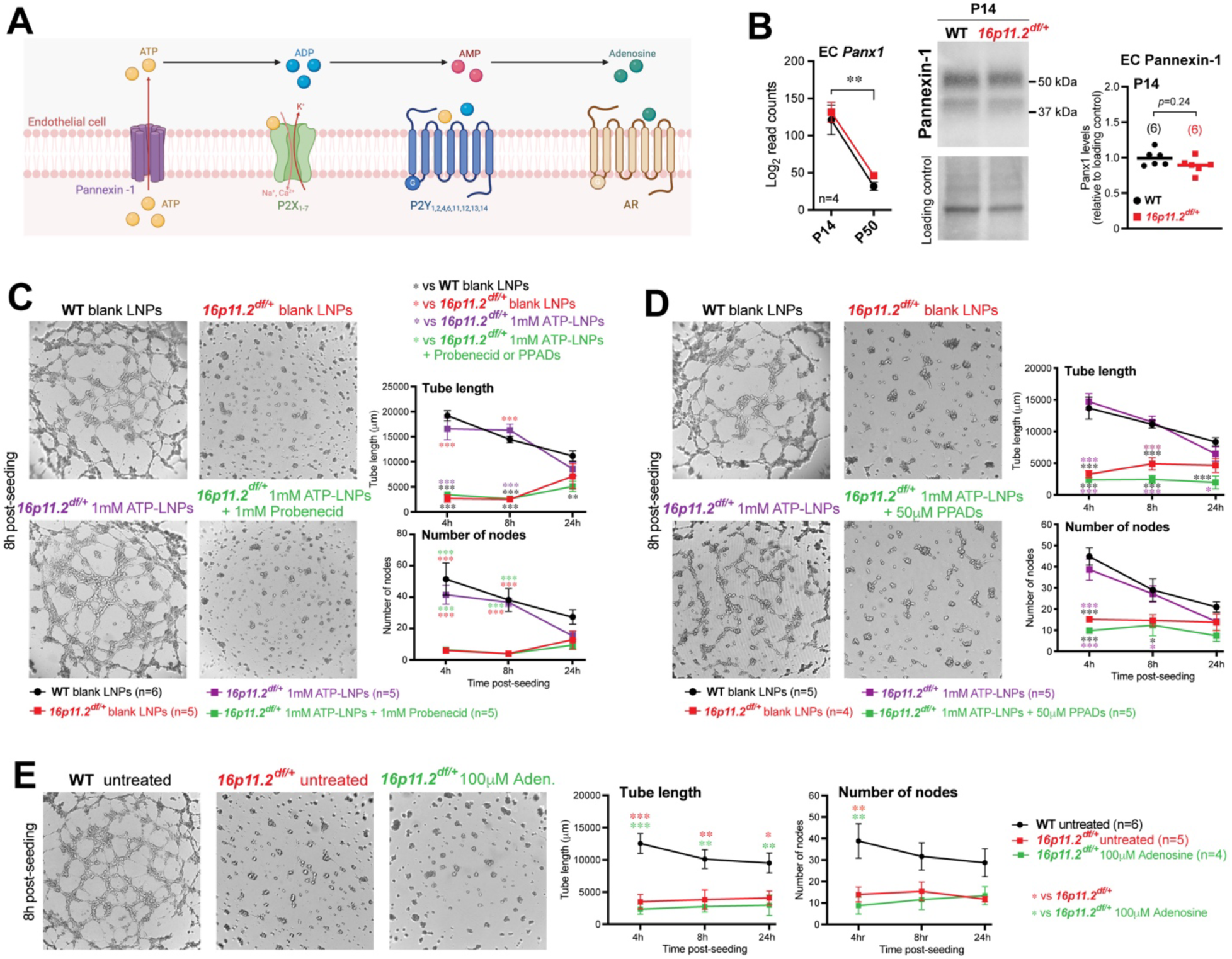
Intracellular ATP supplementation rescues angiogenic activity in 16p11.2-deficient primary brain endothelial cells via Pannexin-1 channels and P2-class purinergic receptors. (A) Schematic representation of ATP release by Pannexin-1 and activation of purinergic signaling AR=adenosine receptor. (B) Left: Expression level of the gene encoding Panx1 in WT and 16p11.2-deficient ECs at P14 and P50 from our previously generated RNAseq database (Ouellette et al., 2020). Data represents mean ± s.e.m. Middle and right: Western blot quantification of relative level of Pannexin-1 protein in 16p11.2-deficient (n=6 mice) and WT (n=6 mice) primary brain ECs. Data represents mean with individual values. \*\**p*<0.01, two-way repeated measure ANOVA and Tukey’s multicomparison *post hoc* test. (C) Probenecid administration prevented rescue of 16p11.2-deficient EC angiogenic activity by intracellular delivery of ATP. Left: representative images of capillary-like networks at the 8h post-seeding time point. Right: quantifications of network densities (total endothelial tube length) and network nodes (total number of branching hubs). LNPs= Lipid nanoparticles. (D) Administration of non-selective P2 receptor antagonist (PPADs) prevented rescue of 16p11.2-deficient EC angiogenic activity by intracellular delivery of ATP. Left: representative images of capillary-like networks at the 8h post-seeding time point. Right: quantifications of network densities (total endothelial tube length) and network nodes (total number of branching hubs). LNPs= Lipid nanoparticles. (E) Adenosine supplementation does not improve 16p11.2-deficient primary brain ECs network formation *in vitro*. Left: representative images of capillary-like networks at the 8h post-seeding time point. Right: quantifications of network tube length and network nodes (total number of branching hubs). untreat.=untreated; Aden.=Adenosine. (B-E) All data shown are from ECs isolated from male mice (n=4-6 animals per group). (C-E) Data are mean ± s.e.m. \**p*<0.05, \*\**p*<0.01, \*\*\**p*<0.001 two-way repeated measure ANOVA and Sidak’s multicomparison *post hoc* test.

Furthermore, brain ECs are known to express various purinergic receptors that are activated by ATP, namely most ionotropic P2X isoforms and several metabotropic P2Y isoforms, all of which take part in maintaining vascular function^27,29^. Considering this, we sought to test if rescue of endothelial function by intracellular ATP is indeed P2 purinergic receptor-mediated. ECs were thus treated with a non-selective P2 purinergic antagonist (pyridoxalphosphate-6-azophenyl-2’,4’-disulfonic acid, PPADS) for 10 minutes prior to each 1mM ATP-LNP administration (*i.e.,* each culture medium change). Administration of 50μM of PPADs abolished the ATP-LNP-mediated rescue in *16p11.2^df/+^* ECs (Figure 2D). These findings demonstrate that LNP-delivered intracellular ATP can indeed exit ECs through Panx1 channels to then act on P2 receptors to mediate functional rescue.

As ATP is rapidly hydrolyzed to adenosine, we next tested whether this rescue was ATP specific. Interestingly, adenosine administration failed to influence endothelial network growth (Figure 2E). In light of this, we focused on extracellular ATP, and whether it can rescue the angiogenic capacity of 16p11.2-deficient ECs *in vitro*. ECs were treated with exogenous ATP at 1μM, 10 μM and 100 μM from cell isolation for 7 days, as previously described (Figure 3A). We found that extracellular ATP supplementation was able to rescue capillary network formation, in a dose-dependent manner (100μM shown in Figure 3A; 1μM and 10μM shown in Figure S4). Lastly, administration of 50μM PPADs abolished the extracellular ATP-mediated rescue in *16p11.2^df/+^* ECs (Figure 3B). At a concentration of 10μM or 100μM, PPADs was less efficient at preventing ATP-mediated rescue (Figures S5A-C). These results confirm that extracellular ATP acts on purinergic P2 receptors to rescue the angiogenic ability of 16p11.2-deficient brain ECs. Overall, these findings indicate that dysfunction of 16p11.2-deficient ECs might result from lack of ATP signaling, rather than lack of ATP itself.

**Figure 3.**
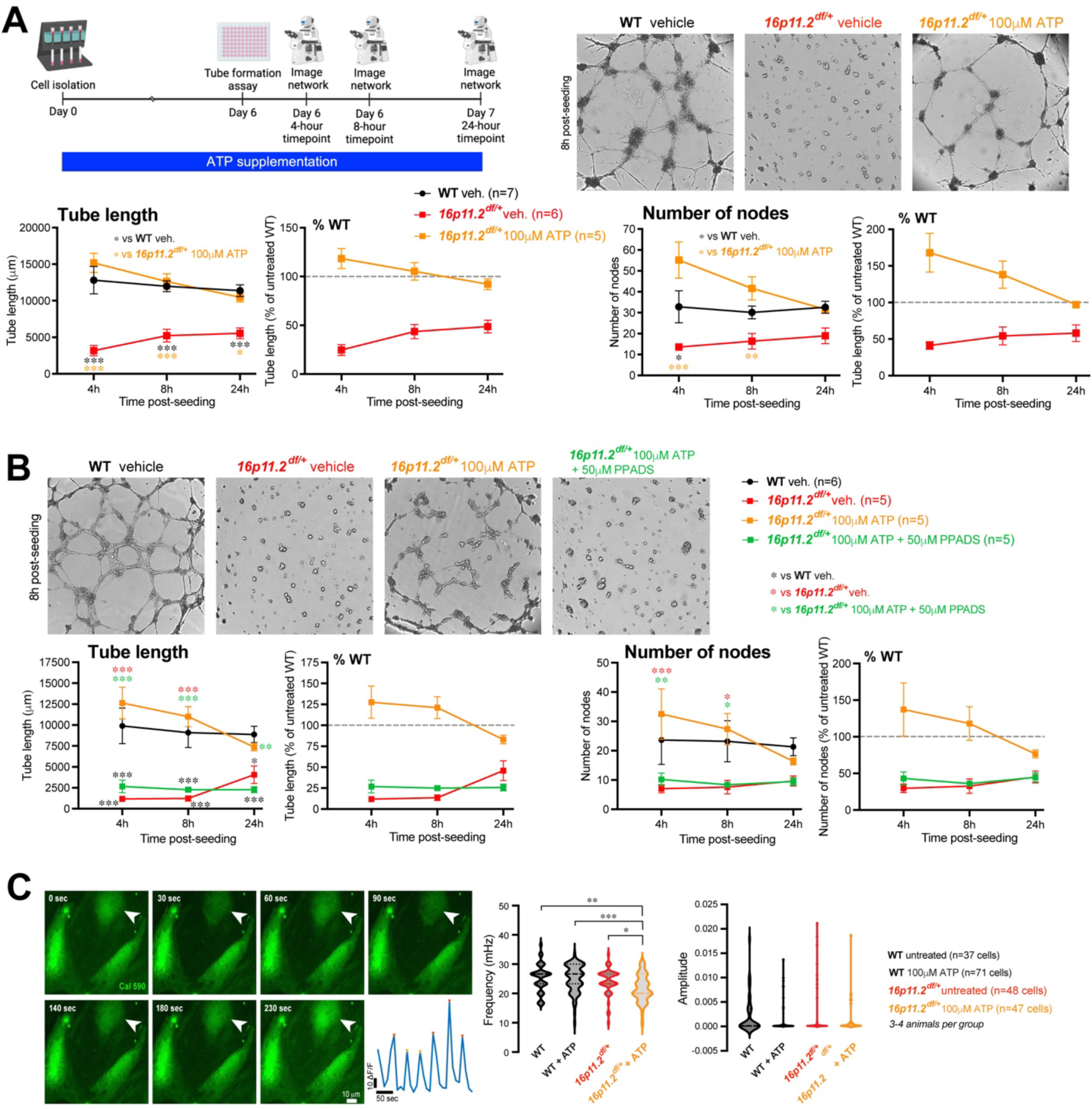
Extracellular ATP-induced rescue is mediated by P2-class purinergic receptors and alters calcium signaling in 16p11.2-deficient ECs. (A) Extracellular ATP rescues 16p11.2-deficient network formation ability. Top left: experimental timeline for treatment of 16p11.2-deficient ECs with extracellular ATP following cell isolation and during a 24-hour *in vitro* network formation assay. Top right: representative images at the 8h post-seeding time point of WT and 16p11.2-deficient untreated ECs and 16p11.2-deficient ECs treated with 100µM of extracellular ATP. Bottom: quantifications of network densities (total endothelial tube length) and network nodes (total number of branching hubs) with representative percent of WT panels. (B) Non-selective P2 receptor antagonist (PPADs) prevents rescue of 16p11.2-deficient EC network formation via extracellular ATP. Top: representative images at the 8h post-seeding time point. Bottom: quantifications of network densities (total endothelial tube length) and network nodes (total number of branching hubs) with representative percent of WT panels. (C) Left: example of time-lapse Ca^2+^ imaging in 16p11.2-deficient ECs and dF/F trace showing Ca^2+^ transients for the cell labeled with white arrowhead. Right: quantification of the frequency and amplitude of Ca^2+^ transients in 16p11.2 deficient and WT ECs. Note, larger decrease in the frequency of Ca^2+^ transients in 16p11.2-deficient, as compared to WT ECs. Violin plots, center line indicating median, n=3-4 animals per group. \**p*<0.05, \*\**p*<0.01, \*\*\**p*<0.001, one-way ANOVA and Tukey multicomparison *post hoc*. (A-C) All data shown are from ECs isolated from P14 male mice. (A and B) Data are mean ± s.e.m or dotted line represents 100 % of WT values (n=5-7 animals per group). \**p*<0.05, \*\**p*<0.01, \*\*\**p*<0.001, two-way repeated measure ANOVA and Sidak’s multicomparison *post hoc* test.

### Altered Ca^2+^-mediated responses to ATP in 16p11.2-deficient brain ECs

Purinergic receptor signaling relies on G protein-coupled receptors resulting in IP_3_-mediated Ca^2+^release from internal stores (P2Y class)^30^ as well as Ca^2+^ permeable ligand-gated ion channels (P2X class)^31^ to establish its function. To gain further insight into ATP-mediated cellular responses, we measured both steady-state intracellular Ca^2+^ levels and Ca^2+^ transients. To assess the steady-state Ca^2+^ level, we used Twitch-2B, a FRET-based ratiometric Ca^2+^ indicator. Our analysis revealed similar steady-state intracellular Ca^2+^ levels under baseline conditions in non-treated WT and *16p11.2^df/+^* ECs (Figure S5D). Interestingly, while ATP supplementation significantly increased intracellular steady-state Ca^2+^ level in WT ECs as compared to WT ECs without ATP, no difference was observed between *16p11.2^df/+^* ECs and *16p11.2^df/+^*with ATP supplementation (Figure S5D). We next loaded ECs derived from WT and *16p11.2^df/+^* mice with Cal590-AM organic Ca^2+^ indicator to record Ca^2+^ transients (Figure 3C). No difference in the frequency and amplitude of Ca^2+^ transients were observed under baseline conditions, in non-treated WT and *16p11.2^df/+^* ECs (Figure 3C). While supplementation with ATP decreased the Ca^2+^ transient frequency in both WT and *16p11.2^df/+^* ECs, the effect was larger in *16p11.2^df/+^* ECs, revealing statistically significant differences in WT + ATP vs. *16p11.2^df/+^* + ATP conditions (Figure 3C). Altogether, our data suggest distinct Ca^2+^ signatures in WT and *16p11.2^df/+^* ECs in response to extracellular ATP supplementation.

### Activation of P2Y2 receptors rescues 16p11.2 deletion phenotypes

Following consultation of our previously published RNAseq database^6^, messenger RNAs (mRNAs) encoding P2-class receptors were found at high expression levels in both genotypes, including *P2rx1*, *P2rx4*, *P2rx7* and *P2ry14* (Figure 4A). The mRNA encoding P2Y2 (*P2ry2)*, a metabotropic receptor important for angiogenesis^29^, was found at relatively low expression levels in WT mice, however its expression nearly doubled in 16p11.2-deficient brain ECs (Figure 4B). This finding being consistent with elevated *P2ry2* expression in plasma of children with ASD^32^. A slight, albeit non-significant, increase in P2Y2 protein levels was seen in 16p11.2-deficient ECs (Figure 4C). Considering this, we decided to engage P2Y2 to test its ability to modulate endothelial function in 16p11.2-deficient ECs. We confirmed localization of P2Y2 receptors at the vessel luminal membrane in P14 brains (Figure 4E). We proceeded to treat 16p11.2-deficient ECs with a selective P2Y2 agonist, Diquafosol tetrasodium (DQS), a compound with efficacy in treating Dry Eye Syndrome post-cataract surgery^33,34^ and clinically-approved in Japan and South Korea (Diquas®). Activation of P2Y2 receptors with 100μM of DQS rescued capillary network formation in *16p11.2^df/+^* ECs (Figure 4D). In light of this and our previous work showing an *ex vivo* endothelium-dependent vasodilation deficit in adult *16p11.2^df/+^* mice^6^, we next sought to determine if P2Y2 engagement will restore *ex vivo* vessel relaxation response of adult *16p11.2^df/+^* cortical parenchymal arterioles (PAs) to adenosine. Following a 30-minute bath perfusion of 100μM of DQS, we found that vessel relaxation of *16p11.2^df/+^* PAs in response to adenosine returned to WT values while *16p11.2^df/+^* arterioles treated with artificial cerebrospinal fluid (aCSF) had a persistent impaired response (Figure 4F). Considering that adenosine evokes a vasodilatory response through an eNOS-dependent mechanism, we assessed eNOS activity in 16p11.2-deficient ECs with or without DQS treatment. We reveal decreased eNOS activity in 16p11.2-deficient ECs compared to WT ECs (Figure 4G), this finding being consistent with the lack of response of 16p11.2-deficient PAs to adenosine (Figure 4F) as well as our previous findings^6^. Interestingly, eNOS activity was rescued in 16p11.2-deficient ECs treated with DQS (Figure 4G) suggesting that P2Y2 receptor activation increases eNOS activity to enable vasodilation of 16p11.2-deficient PAs in response to adenosine.

**Figure 4.**
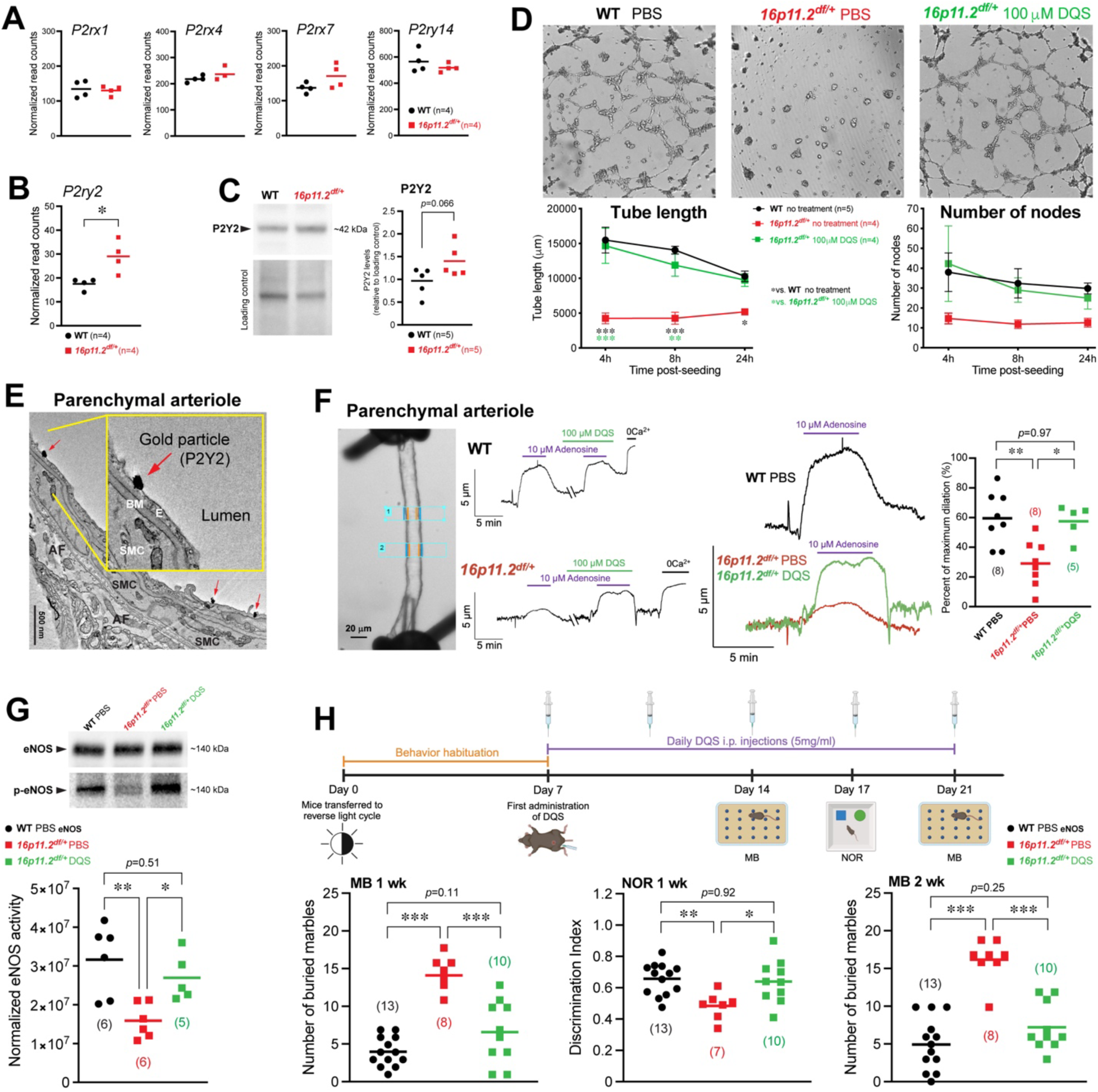
Pharmacological P2Y2 receptor activation rescues 16p11.2-deficient endothelial function and normalizes adult *16p11.2^df/+^*mouse behavior. (A) Expression level of genes encoding P2-class receptors in WT and 16p11.2-deficient ECs from our previously generated RNAseq database (Ouellette et al., 2020). (B) A higher expression level of the gene encoding P2Y2 receptors was identified in 16p11.2-deficient ECs. \**p*<0.05, two-tailed Mann-Whitney test. (C) Western blot quantification of relative P2Y2 protein levels in primary mouse brain ECs (n=5) isolated from 16p11.2-deficient and WT littermates. *p*=0.066, two-tailed Mann-Whitney test. (D) Top: representative images at the 8h post-seeding time point of the *in vitro* network formation assay following treatment of ECs with P2Y2 agonist Diquafosol tetrasodium (DQS). Bottom: quantification of network densities (total endothelial tube length) and network nodes (total number of branching hubs). Data represents mean ± s.e.m (n=4-5 animals per group). \**p*<0.05, \*\**p*<0.01, \*\*\**P*<0.001, two-way repeated measure ANOVA and Sidak’s multicomparison *post hoc*. (E) Electron micrograph of immunogold labeling on a parenchymal arteriole showing localization of P2Y2 receptors at the luminal membrane (red arrows). AF, astrocytic endfoot; BM, basement membrane; E, endothelium; SMC, smooth muscle cell. (F) Vascular reactivity of cortical parenchymal arterioles (PAs) *ex vivo* during DQS treatment. Changes in arteriole diameter were measured in response to adenosine. Representative image of PA preparations (left) and traces of arteriolar diameter of WT (top middle) and *16p11.2^df/+^* (bottom middle) PA during DQS treatment. Right: quantification of *ex vivo* vascular reactivity following DQS treatment revealed a rescue in vasodilatory responses of *16p11.2^df/+^*arterioles. Data represents n=5-8 animals per group. (G) Western blot quantification of eNOS activity from 16p11.2-deficient ECs treated with DQS (n=5), 16p11.2-deficient ECs (n=6) and WT ECs (n=6). (H) Top: schematic representation of DQS injections and behavioral timeline. Bottom: behavioral assessments of adult *16p11.2^df/+^* males at P50 demonstrated reduced marble burying (MB) following a one-week (left) and a two-week DQS treatment (right) compared with PBS treated *16p11.2^df/+^* males. Bottom middle: improved recognition memory in *16p11.2^df/+^* males treated with DQS following a novel-object recognition task (NOR) compared to PBS treated *16p11.2^df/+^* males. Data represents n=7-13 animals per group. (A-D) Data shown are from ECs isolated from P14 male mice. (A-C, F, G and H) Data represents mean with individual values. (F-H) \**p*<0.05, \*\**p*<0.01, \*\*\**p*<0.001, one-way ANOVA and Tukey multicomparison *post hoc*.

Prior work has demonstrated that *16p11.2^df/+^* mice are characterized with repetitive movements and impaired novel-seeking behaviors presented as novel-object recognition deficits, behaviors frequently found in 16p11.2 deletion carriers^6,35,36^. To assess the extent of DQS treatment rescue on 16p11.2 deletion phenotypes, we next tested if DQS administration can rescue 16p11.2 deletion associated-behaviors. Adult *16p11.2^df/+^* mice were intraperitoneally (i.p.) injected with 5mg/ml of DQS at 1% of body weight once a day for a total of two weeks (one week prior to behavior assays and one week during behavior assays) (Figure 4H). Notably, daily injections of DQS did not impact survival (Figure S6E). We observed that adult *16p11.2^df/+^* male mice treated with DQS exhibited reduced marble burying, indicative of reduced repetitive behaviors, following one week of treatment compared to PBS-treated *16p11.2^df/+^* mice, which was maintained after two weeks of treatment (Figure 4H). Moreover, DQS treatment of *16p11.2^df/+^*mice also rescued recognition memory (*i.e.* cognitive deficits) during a 2-day novel-object recognition task, returning values to WT levels, compared to PBS-treated *16p11.2^df/+^* mice (Figure 4H). Thus, suggesting that P2Y2 receptor activation may represent a strategy to maintain brain EC function and rescue ASD-related behaviors in 16p11.2 deletion ASD syndrome.

## Discussion

We demonstrate that endothelial dysfunction caused by 16p11.2 haplodeficiency is associated with a bioenergetic failure in ECs. Our work also reveals that purinergic signaling activation via P2Y2 receptor engagement rescues 16p11.2 deletion-associated endothelial phenotypes and adult animal behavior. These findings highlight metabolic reprogramming of brain ECs as a potential therapeutic avenue for ASD.

Our investigation into 16p11.2-deficient ECs metabolism revealed a significant reduction in intracellular metabolites. While metabolomics studies have been performed in the context of ASD^37–39^, this work provides the first metabolomics approach focused on brain ECs in ASD, unraveling endothelial-specific metabolic changes that contribute to 16p11.2 deletion-related dysfunctions. Metabolite profiling of 16p11.2-deficient ECs helped us identify a reduction of high-energy molecules including ATP, a reduced adenylate energy charge, and a reduced ATP-to-ADP ratio, altogether indicative of a lack of energy availability.

While here we do not find altered mitochondrial function 16p11.2-deficient ECs, altered mitochondrial parameters are emerging contributors to ASD^40^. We do find reduced mitochondrial mass *in vitro* using ECs isolated from 14-day old mice, reiterating our findings in 50-day old 16p11.2-deficient mice *in vivo*^7^. This suggest that altered mitochondria parameters in development might have an early impact on brain EC metabolism. However, we do not provide causal evidence supporting such link. Moreover, reduced mitochondrial density in P14 16p11.2-deficient ECs could not be attributed to altered mitochondrial dynamics or altered biogenesis. This suggests that other mechanisms might be at play to downregulate mitochondrial biogenesis postnatally, such as increased mitophagy. Although the precise connection between 16p11.2 hemideletion and reduced mitochondrial density is not known at this time, mitochondrial alterations represent a valid candidate to explain functional disturbances in a range of physiological systems.

Impact of the lack of bioavailable energy was particularly evident when brain ECs were challenged or, in other words, when a functional task was required (*i.e.* angiogenesis, vascular reactivity)^6^. It has been recently shown that ATP is essential for endothelial control of vascular tone. A recent study showed that when mitochondrial ATP production was blocked *in vivo* or *ex vivo*, ECs did not respond to vasodilator acetylcholine in both cerebral and mesenteric arteries^21^. This is in line with our prior work demonstrating that both middle cerebral and mesenteric arteries from 16p11.2-deficient mice respond poorly to acetylcholine^6^. Thus, decreased intracellular ATP in 16p11.2-deficient brain ECs suggests that 16p11.2 deletion-associated bioenergetic failure may be responsible for endothelial dysfunction. While the reduced mitochondrial density in 16p11.2-deficent ECs was slight, albeit significant, recent evidence has shown that even a small dose of one mitochondrion, following mitochondrial transplantation, can significantly increase ATP levels in cells^41^, highlighting the possible impact of the minor changes in mitochondrial mass on intracellular ATP levels.

Strikingly, increasing ATP bioavailability intra- or extra-cellularly rescued 16p11.2-deficient EC function, further suggesting that lack of ATP might cause deficits. Extracellular ATP has emerged as a signaling regulator in several cellular processes including proliferation, angiogenesis, and vessel homeostasis^21^. Brain ECs release ATP in response to various stimuli, triggering signaling events via ligand gated (P2X) and/or G-protein-coupled (P2Y) purinergic receptors^42,43^. In line with ATP regulating key vascular features via activation of purinergic receptors, we provide novel evidence that ATP-mediated rescue of 16p11.2-related endothelial dysfunction relies on P2-class receptors expressed by ECs. Indeed, non-selective pharmacological blockade of P2 receptors abolished the intra- and extra-cellular ATP-mediated rescue. We also reveal that intracellular ATP exits ECs via Pannexin-1 channels to subsequently act on membrane-bound P2 receptors, a recently established EC mechanism^27^.

P2-class purinergic receptor activation via ATP can modulate Ca^2+^ responses. Interestingly, we observed differences in Ca^2+^ transients in 16p11.2-deficient ECs, as compared to WT ECs, in response to extracellular ATP. Pannexin-1 channels are activated by increases in intracellular calcium leading to the release of ATP from the cell and so ATP acting on P2 receptors^27^. While it is yet unclear how these distinct Ca^2+^ responses to ATP may underlie the rescue of 16p11.2-deficient ECs, our data hint on Ca^2+^ as a possible signaling molecule mechanistically linking purinergic receptor activation to intracellular pathways leading to restoration of angiogenic ability of 16p11.2-deficient ECs.

We specifically highlight the involvement of P2Y2 receptor activation as a mechanism to rescue 16p11.2-related endothelial dysfunction. P2Y2 receptors have a prominent role in angiogenesis, as their activation promotes endothelial sprouting and vascular tube formation^29^. Inhibition of P2Y2 with a selective antagonist reduced tube length in human umbilical vein endothelial cells (HUVECs)^29^. These findings emphasize that lack of P2Y2 activation in 16p11.2-deficient ECs due to decreased ATP availability might be sufficient to drive functional changes.

Furthermore, P2Y2 receptors have been associated with vascular function^29^. Mice with endothelial P2Y2-knockout displayed dysregulated vasodilatory responses during increased blood flow^44^. Here, we show that P2Y2 agonism reinstates parenchymal arteriolar vascular reactivity *ex vivo* in 16p11.2-deficient mice via increased eNOS activation, thus further supporting the involvement of P2Y2 in vascular function. Moreover, systemic P2Y2 activation rescued 16p11.2 deletion-associated behaviors highlighting that lack of purinergic signaling is involved in 16p11.2 deletion phenotypes, and the possibility of modulating purinergic signaling in ASD. While we confirmed that endothelial functional rescue was specifically conferred by ATP and not by adenosine, recent evidence has associated adenosine with improved social novelty behaviors in adult female *16p11.2^df/+^*mice^45^. As all our experiments were perform using male mice, it is possible that these beneficial effects of adenosine in female mice behavior are independent of vascular features.

While the purinergic system has been linked to ASD, it is in fact anti-purinergic therapies that led to improvements of autism-like behavior in rodents^39,46^. In certain instances, elevated extracellular ATP in the blood of ASD patients have been associated with metabolic changes and suggested to activate the cellular danger response (CDR), leading to detrimental stress responses (*i.e*. changes in mitochondrial function, and innate immune activation)^39^. As we demonstrate that ATP exits the cells to act on adjacent P2 receptors, the CDR could explain the lack of rescue following delivery of a higher dose of intracellular ATP (*i.e.* 5mM ATP-LNPs). As such, higher extracellular ATP may negatively impact EC function. Along these lines, extracellular ATP/adenosine imbalance has been linked with seizures, which are common in ASD patients^47^. In addition, maintained extracellular ATP levels using non-hydrolysable ATP, an analogue which cannot be converted to adenosine, has been shown to have lethal effects in mice^48^. These findings not only support the involvement of purinergic signaling in ASD but also show that a purinergic signaling threshold should be considered when designing these therapies.

Here, we emphasize the importance of studying ASD with a holistic lens, as these disorders remain predominantly considered a result of neuronal malfunction. Revealing new disease hallmarks specific to endothelial metabolism should help broaden therapeutic strategies for ASD.

## ACKNOWLEDGMENTS

This research was funded by grants from the Canadian Institutes of Health Research (#388805, #506513) to B.L., and a Scholarship from the Canadian Institutes of Health Research to J.O.

The following patent application related to this study has been filed:

## NEURODEVELOPMENTAL DISORDER TREATMENT

Patent Cooperation

Treaty Application No. PCT/CA2025/050486, filed 2025/04/03. *Patent Status: Pending*

## METHODS

### Animals

All animal procedures were approved by the University of Ottawa Animal Care Committee and conducted in accordance with guidelines of the Canadian Council on Animal Care. *Mouse husbandry.* All mice were bred in house and housed maximum five per cage with free access to water and food. Males *16p11.2^df/+^*(Jackson laboratory, stock #013128; mixed B6/129 background) were crossed with WT females of the same background (Jackson laboratory, stock #101043) to obtain hemizygous *16p11.2^df/+^* offspring as well as WT littermates. As recommended by Jackson Laboratory to improve *16p11.2^df/+^* pup survival, breeding cages were supplemented with breeding chow (#2019, Envigo Teklad) and DietGel (#76A, ClearH_2_O) up to weaning age. All *in vitro* experiments were conducted using P14 mice while ex vivo and in vivo experiments were conducted with 7-9 weeks old mice.

*Genotyping. 16p11.2^df/+^* mice and WT littermates were genotyped using the following primers: 5′-CCTCATGGACTAATTATGGAC-3′ (forward) and 5′-CCAGTTTCACTAATGACACA-3′ (reverse), with a PCR product of 2.2 kb for *16p11.2^df/+^* mice ^49^.

### Primary mouse brain EC isolation

All mice were euthanized by cervical dislocation. The cerebral cortex was dissected in cold HBSS without calcium and magnesium using autoclaved tools submerged in 100% ethanol for 30 minutes prior dissections. The cortex was minced in 2-3 mm pieces and dissociated in Neural Tissue Dissociation Kit P compounds (Miltenyi Biotec, 130-092-628) to obtain a cell suspension. Cell isolation procedures were completed according to the manufacturer’s instructions. The cell suspension was incubated with CD31-coated magnetic microbeads (Miltenyi Biotec, 130-097-418) and placed on a magnetic MACs separator to isolate ECs. ECs isolated from one mouse were seeded in a plate coated with attachment factor protein 1X (ThermoFisher Scientific, S006100). For a pure EC population, ECs were cultured in an EC specific medium (Lonza, CC-3202) which was replaced 48 hours post-seeding and every 48 hours until appropriate level of confluency was reached for subsequent processing (40-60% for immunocytochemistry; 90-100% for metabolomics, protein extraction, tube formation and 80% for calcium experiments).

### Immunocytochemistry

Primary ECs were cultured until 40–60% confluence (3 days post-isolation) on glass coverslip for microscopy analysis. Before staining, ECs were washed twice in pre-warmed PBS and fixed for 10 min in cold 4% PFA. Fixed cells were permeabilized in a blocking solution containing 10% donkey serum, 0.1% PBT and 0.5% cold water fish skin gelatin. The primary antibodies anti-CD31 (1:200, BD Pharmingen) and anti-TOM20 (1:2000, Proteintech) were diluted in blocking solution and incubated for 2 hours at room temperature. Cells were then rinsed with 0.2% PBT. AlexaFluor species-specific secondary antibodies were diluted in blocking solution at 1:300 and incubated with cells for 60 minutes. Cells were washed with 0.2% PBT then 0.1 M PB. Glass coverslips, with stained ECs, were mounted on slides using Fluoromount G and imaged (40x objective) with a Zeiss Axio Imager M2 microscope equipped with a digital camera and ApoTome.2 module. Images consisted of 4uM thick z-stacks (0.250µM/slice) in which 5-10 ECs were visible per image, a total of 10 images were taken per animal. To quantify TOM20 and CD31 immunostaining, 40X ApoTome images (16-bit Tiff files) were processed using custom scripts written in the Python programming language. Mitochondria were detected using scale-space maxima of the Laplacian of Gaussian (LoG) operator ^50^. Specifically, the images were first convolved with four LoG filters with standard deviations of 1.0, 1.33, 1.66, and 2.0 pixels. Peaks having a value larger than 0.03 on the scale-space representation were considered to represent mitochondria. ECs nuclei were also automatically identified to avoid mitochondria detection inside the nuclei. Pixels having an intensity smaller than 10 on nuclei (DAPI) images were considered to belong to the background. Connected components with less than 75 pixels were removed, and remaining holes were filled. The obtained connected components were associated with nuclei of the ECs. Mitochondria detected inside the identified nuclei were then removed by masking. ECs within each image were manually outlined and mitochondria density per cell was calculated as the number of mitochondria divided by cell area. The values were then averaged among the outlined cells. To calculate fragmentation, a mitochondria neighborhood graph was created to identify their spatial relationship. To do so, the highest affinity paths between all pairs of mitochondria having a distance smaller than 15 pixels between them were identified. The affinity of a path consists of the sum of the image intensities of a connected sequence of pixels between two mitochondria. The lowest intensity value along the highest affinity path between two mitochondria defined their affinity. Intuitively, if there is a connected sequence of pixels between two mitochondria such that no pixel in the sequence has a low-intensity value, the two mitochondria likely belong to a single, non-fragmented, structure. A graph having mitochondria as nodes was constructed, two nodes were connected if the affinity between the respective mitochondria was larger than 0.25 times the average intensity of the two mitochondria. The fragmentation was then calculated as the number of connected components in the graph divided by the number of mitochondria.

### Western blots

#### Protein extraction

For endothelial proteins, primary brain endothelial cells were isolated and cultured as previously described ^7,51^. To lyse cells, 50µL of RIPA buffer (NaCl 150mM, Sodium Deoxycholate 0.5%, SDS 1%, NP-40 1% in Tris 50mM pH 8.0) with protease and phosphatase inhibitors was added to each well and left on ice for 30 min. The cells were scraped off the culture dish and transferred to a 1.5mL tube then sonicated for 1 minute on ice. Tubes were stored at - 20°C overnight and then centrifuged at 13,000rpm for 10 min. The protein concentration of the collected supernatant was quantified using Pierce BCA Protein assay (kit #23227, ThermoScientific, Illinois, USA).

#### Protein extraction for AMPK samples

For endothelial proteins, primary brain ECs were isolated and cultured as previously described. To lyse cells, 100µL of AMPK extraction buffer (50 mM Tris-HCL, pH 7.5, 1 mM EDTA, 1 mM EGTA, 0,27M sucrose, 1% Triton X-100 and 10mM DTT with protease and phosphatase inhibitors) was added to each well. The bottom of the well was scraped for one minute. The supernatant was transferred to 1.5mL tubes and then sonicated, incubated on ice for 15 minutes and centrifuged at 13 000 rpm for 15 minutes at 4 °C to remove cell debris. The protein concentration of the collected supernatant was quantified using Pierce BCA Protein assay (kit #23227, ThermoScientific, Illinois, USA).

#### Immunoblotting

Twelve (12) µg of protein (MFN1, MFN2, OPA1 and DRP1), forty (40) µg of protein (AMPK and phospho-AMPK) or fifteen (15) ug of protein (eNOS and p-eNOS) were loaded into the wells of SDS-acrylamide gels (GX Stain-free FastCast 12% Biorad gels, Bio-Rad, ON, CAN, #1610184) and separated by a constant current of 100 V for 120 min in running buffer (SDS 35mM, Tris 250mM, Glycine 1865mM). After UV activation of the gels, the proteins were then transferred to a nitrocellulose membrane in ice-cold transfer buffer (Tris 48mM, Glycine 38mM, methanol 20% v/v) for 30 min at 100 V. After the transfer, the nitrocellulose membranes and gels were imaged to quantify the total protein transferred. The membranes were then blocked with 5% skim milk in 1M PBS (or 5% BSA in TBST for AMPK and p-eNOS samples) for 1 h at room temperature and incubated with primary antibodies raised against either AMPK (1:1000, Cell Signaling, 2532), phospho-AMPK (1:800, Cell Signaling, 2535), DRP1 (1:2000, BD Biosciences, 611113), MFN1 (1:1000, Abcam, ab126575), MFN2 (1:1000, Abcam, ab56889), OPA1 (1:2000, Abcam, ab42364), PGC-1a (1:1500, Millipore, ST1202), eNOS (1:1000, Cell Signaling, 32037), p-eNOS (1:1000, Cell Signaling, 9571) in TBST (Tris 50mM, NaCl 150mM, Tween 20 1% v/v) overnight at 4°C. The membranes were then washed with TBST (3 x 10 min) and incubated at room temperature for 1 h with the secondary antibody (1/10,000, Fisher Scientific, #PR-W4011), also diluted in TBST. After a final wash (TBST, 3 x 10 min), the proteins were detected by enhanced chemiluminescence (ECL; 1⁄4, Fisher Scientific, #34579) and imaged with the Odyssey Imaging system. The samples were normalized to the total protein content (AMPK) and to a loading control (OPA1, DRP1, MFN1&2, PGC1a) and calculated as relative to group average.

#### Immunoblotting for Pannexin-1

Thirty (30) μg protein was separated on a 10% SDS-PAGE gel according to manufacturer’s protocol (TGX Stain-Free FastCast, BioRad) and transferred to polyvinylidene difluoride (PVDF) membranes. Membranes were processed according to manufacturer’s protocol (TGX Stain-Free FastCast, BioRad) and incubated overnight at 4°C with primary antibody anti-Pannexin-1 (1:500, Cell Signaling, Cat# 91137) in Tris-buffered saline (Tris 50 mM and NaCl 150 mM, pH 8.0) containing 0.05% Tween (TBST) in 1% non-fat dry milk. The membranes were washed and incubated with horseradish peroxidase–conjugated secondary antibody: Anti-Rabbit HRP or Anti-mouse HRP (Promega or Thermo Fisher Scientific) 1:2000 in TBST containing 5% non-fat dry milk. Immunoreactive proteins were visualized by chemiluminescent solution (SuperSignal West Dura; Pierce Biotechnology). Densitometric, semiquantitative analysis of Western blots was performed using Image Lab software and normalized to total protein.

#### Targeted metabolomics

Primary ECs were split once and cultured until 90-100% confluency was reached. ECs from 3 mice were pooled to have at minimum 1.5million cells per sample (n). Prior sample collection, cells were washed 3 times with 1mL of ammonium formate per well of a 6-well plate. For sample collection, 80uL of ice cold methanol:water (1:1) was added to each well and cells were collected using a cell scraper. The cell slurry was transferred to a pre-chilled 2 mL tube containing 6 washed ceramic beads (1.4 mm) and kept at −80 °C until the day of metabolite extraction.

On the day of extraction, samples were vortexed 10s, followed by adding 220 µL of acetonitrile and vortexed a second time for 10s. Cell lysis was done by bead beating for 60 s at 2000 rpm (bead beating was done twice). Samples were then incubated with a 2:1 dichloromethane:water solution on ice for 10 minutes. The polar and non-polar phases were separated by centrifugation at 4000g for 10 minutes at 1°C. The upper polar phase was dried using a refrigerated CentriVap Vacuum Concentrator at −4°C (LabConco Corporation, Kansas City, MO). Samples were resuspended in water and run on an Agilent 6470A tandem quadruple mass spectrometer equipped with a 1290 Infinity II ultra-high performance LC (Agilent Technologies) utilizing the Metabolomics Dynamic MRM Database and Method (Agilent), which uses an ion-pairing reverse phase chromatography. This method was further optimized for phosphate-containing metabolites with the addition of 5 µM InfinityLab deactivator (Agilent) to mobile phases A and B, which requires decreasing the backflush acetonitrile to 90%. Multiple reaction monitoring (MRM) transitions were optimized using authentic standards and quality control samples. Metabolites were quantified by integrating the area under the curve of each compound using external standard calibration curves with Mass Hunter Quant (Agilent). No corrections for ion suppression or enhancement were performed, as such, uncorrected metabolite concentrations are presented. Multivariate statistical modelling was performed on log-transformed, mean-centered and pareto-scaled data using MetaboAnalyst 5.0 (www.metaboanalyst.ca). Partial least squares discriminant analysis (PLS-DA) was performed on the identified metabolites. The variables highly contributing to the group separation were selected with a variable importance in projection (VIP)≥1. The clustering analysis and pathway analysis were performed as well using MetaboAnalyst 5.0. Statistical analysis was performed by one-way analysis of variance (ANOVA) followed by Tukey’s post hoc test. P values <0.05 were considered statistically significant. Pairwise comparisons were completed to eliminate batch effect. Data for genotype comparison is presented as box and whisker plots and statistical analysis was performed by unpaired t test. The box plot analysis was performed using Prism 10 (GraphPad Software, La Jolla, CA, USA).

#### Seahorse Cell Mito Stress test assay

Agilent Seahorse Cell Mito Stress test XF96 kit was prepared according to the manufacturer’s instructions. 24 hours before the start of the assay, ECs were harvested and seeded at a density of 10,000 cells per well in a Seahorse XFe96 well plate coated with 1X attachment factor. The sensor cartridge was submerged in XFe calibration buffer overnight in a non-CO_2_ incubator at 37°C. On the day of assay, Seahorse XF Base Medium was supplemented with 1mM pyruvate, 10mM glucose and 2mM glutamine. Chemical compounds were reconstituted using assay medium. Cells were washed twice with assay medium and incubated for 1 hour in a non-CO_2_ incubator at 37°C. Reconstituted compounds were loaded onto a sensor cartridge at the following final concentrations: Oligomycin (port A, 1.0μM), FCCP (port B, 1.0μM) and Rotenone/antimycin A (port C, 0.5μM). Sensor cartridge with loaded compounds were placed in a pre-heated Seahorse XFe96 analyzer for calibration. Following the 1 hour incubation and sensor cartridge calibration, the cells are placed in the Seahorse XFe96 analyzer for subsequent mitochondrial function assessment. Once the assay was completed, cells were fixed with 4% PFA, washed twice with 1X PBS and stained with DAPI for cell counting. Cells within each well was counted using a EVOS FL Auto imaging system for normalization of parameters. Assay was completed and analyzed with the Agilent Seahorse Wave Software.

### ATP-LNPs

#### Materials

Adenosine triphosphate (ATP) (150266) was purchased from MP Biomedicals (Solon, OH). Cholesterol (8667) was procured from Sigma-Aldrich (St. Louis, MO). 1,2-Dimyristoyl-rac– glycero-3-methoxypolyethylene glycol-2000 (PEG-DMG) (8801518) and 1, 2-distearoyl-sn-glycero-3-phosphocholine (DSPC) (850365P) were obtained from Avanti Polar Lipids (Alabaster, AL). The ionizable cationic lipidoid, 1,1‘-((2-(4-(2-((2-(bis (2 hydroxy dodecyl) amino) ethyl) (2hydroxydodecyl) amino) ethyl) piperazin1yl) ethyl) azanediyl) bis(dodecan-2-ol) (C12-200) was a generous gift from Dr. Muthiah Manoharan, Alnylam Pharmaceuticals (Cambridge, MA). Alexa Fluor 647-labeled ATP (AF647-ATP) (A22362) was obtained from ThermoFisher Scientific (Waltham, MA). Phosphate-buffered saline (PBS) was procured from Hyclone Laboratories (Logan, UT). All reagents were used as obtained unless stated otherwise.

#### Preparation of ATP-loaded LNPs (ATP-LNPs)

Lipid nanoparticles (LNPs) were prepared using previously reported methods ^22,52^. Briefly, C12-200 (ionizable cationic lipid), cholesterol, PEG-DMG, and DSPC were dissolved in ethanol at a molar ratio of 50/10/38.5/1.5. The representative scheme for the formulation of ATP-LNPs is depicted in **Table 1**. The aqueous phase consisted of an ATP solution at 33.73 mg/mL prepared in 1*x* PBS, pH 7.4. The ethanolic phase was gradually added into the aqueous phase under continuous vortexing for 30 seconds using a Fisher benchtop vortexer knob at position ‘7’. The resulting ATP-LNP formulations were prepared with a final ATP concentration of 40 mM. Alexa fluor 647-labeled ATP-LNPs were prepared by volumetrically spiking unlabeled ATP with Alexa fluor 647-labeled ATP at a ratio of 1:1000 (Alexa fluor 647-labeled ATP: unlabeled ATP).

**Table 1.** Formulation scheme of ATP-LNPs^22^.

| Component | w/w% in LNP mixture | Stock concentration (mg/mL) | Volume of the stock solution (μL) |
| --- | --- | --- | --- |
| **Ethanol phase** |  |  |  |
| C12-200 | 50 | 1 | 12.5 |
| Cholesterol | 38.5 | 1 | 3.1 |
| PEG-DMG | 1.5 | 0.1 | 13.6 |
| DSPC | 10 | 0.1 | 16.9 |
| Ethanol | - | - | 78.9 |
| **Total ethanol phase volume** |  |  | 125 |
| **Aqueous phase** |  |  |  |
| ATP | 40 mM | 33.73 | 300.7 |
| 1x PBS | - | - | 74.3 |
| **Total aqueous phase volume** |  |  | 375 |
| **Final volume of LNPs prepared (μL)** |  |  | 500 |

#### Dynamic Light Scattering

The particle characteristics and colloidal stability of ATP-LNPs were studied by measuring their average particle diameters and dispersity indices using dynamic light scattering on a Malvern Zetasizer Pro (Malvern Panalytical Inc., Westborough, PA). Particle sizes and dispersity indices of ATP-LNPs were measured over 14 days with intermittent storage at 2-8°C. For dynamic light scattering measurements, LNPs were diluted to a final ATP concentration of 40 mM in 1*x* PBS pH 7.4. The data are depicted as the mean value along with the standard deviation (SD) derived from triplicate measurements.

#### Transmission Electron Microscopy

LNPs were prepared for negative stain electron microscopy by depositing a 3 µL volume onto a freshly glow-discharged continuous carbon-coated copper grid and subsequently treating it with a 1% uranyl acetate solution for staining purposes. Grids were introduced into a Thermofisher TF20 electron microscope, outfitted with a field emission gun, and imaged using a TVIPS XF416 CMOS camera to visualize LNP dimensions. The electron microscope was operated at 200 kV and the contrast was enhanced using a 40 µm objective aperture. Electron micrographs were captured at a nominal 19,000*x* magnification using a post-column magnification of 1.3*x*, corresponding to a calibrated pixel size of 5.6 Ångstroms at the sample level. The images were acquired utilizing TVIPS Emplified software in movie mode to correct for drift.

#### ATP-LNPs uptake

Primary brain ECs were cultured until 80% confluency on a cover-glass and treated with AF647-labeled ATP-LNPs or blank-LNPs following cell isolation. Primary ECs were washed twice with pre-warmed PBS and stained with Hoechst (Invitrogen, C10339 G) for 5 minutes. The stain was removed and cells were washed twice with PBS. Cover glasses were mounted on slides and imaged with a Zeiss Axio Imager M2 microscope equipped with a digital camera and ApoTome.2 module.

### In vitro treatment protocols

#### Intracellular ATP supplementation (ATP-LNPs)

Primary ECs were treated with 1mM or 5mM of ATP-LNPs or blank-LNPs following cell isolation and once cells reached 90-100% confluency, they underwent an *in vitro* network-formation assay. ATP-LNPs were directly added to EC specific culture media (Lonza, CC-3202) and was replaced every 48hrs until required confluency was reached.

#### Extracellular ATP supplementation

Primary ECs were treated with 1μM, 10μM or 100μM of ATP (Sigma Aldrich, A7699) following cell isolation and once cells reached 90-100% confluency, underwent an *in vitro* network-formation assay. ATP was directly added to EC specific culture media (Lonza, CC-3202) and was replaced every 48hrs until required confluency was reached.

#### Adenosine supplementation

Primary ECs were treated with 100μM of adenosine (Sigma Aldrich, A4036) or NH_4_OH (vehicle, Sigma Aldrich, 09859) following cell isolation, once cells reached 90-100% confluency, underwent an *in vitro* network-formation assay. Adenosine solution was prepared according to the manufacturers. Briefly, adenosine was dissolved in 1M NH_4_OH using heat (70°C) for 5 mins and maintained at room temperature protected from light. Adenosine was directly added to EC specific culture media (Lonza, CC-3202) and was replaced every 48hrs until required confluency was reached.

#### Non-selective P2 antagonist treatment

Primary ECs were treated with 10μM, 50μM or 100μM of pyridoxalphosphate-6-azophenyl-2’,4’-disulfonic acid (PPADs) tetrasodium salt (Tocris, 0625) for 10 minutes prior to addition of 100μM of ATP and replaced every 48hours until cells reached 90-100% confluency. Once confluency was reached, ECs underwent an *in vitro* network-formation assay.

#### Non-selective P2 antagonist (PPADs) and ATP-LNP treatment

Primary ECs were treated with 50μM of PPADs for 10 minutes prior administration of 1mM of ATP-LNPs. Cell culture media was replaced every 48 hours until cells reached 90-100% confluency. Once confluency was reached, ECs underwent an *in vitro* network-formation assay.

#### P2Y2 agonist (diquafosol tetrasodium, DQS) treatment

Primary ECs were treated with 100μM of DQS (Sigma Aldrich, SML3058) following cell isolation, throughout culturing and during the tube formation assay. DQS was directly added to EC specific culture media (Lonza, CC-3202) and was replaced every 48hrs until required confluency was reached (90-100%).

#### In vitro network-formation assay

EC network-formation assays were performed using growth-factor-reduced Matrigel (BD Bioscience, cat. no. C354230). Growth-factor-reduced condition includes: EGF < 0.5 ng ml^−1^, PDGF < 5 pg ml^−1^, IGF1 = 5 ng ml^−1^ and TGF-β = 1.7 ng ml^−1^. In brief, each well of a 96-well plate was coated with 50 μl of Matrigel and incubated at 37 °C for 30 min to promote polymerization. ECs were collected with TryplE (Gibco, 12604013) and counted. A total of 2 × 10^4^ cells were seeded in each well with 150 μl EC specific media (Lonza, CC-3202) or EC media supplemented with ATP-LNPs, ATP, Adenosine, ATP+PPADs, ATP-LNPs+ PPADs or DQS. TIFF images of capillary-like networks were captured using a Zeiss Axio Image M2 microscope equipped with a digital camera at 4, 8, and 24h post-seeding (mouse ECs). Images were processed using the Angiogenesis function of ImageJ ^53^.

#### Steady-state cytosolic Ca^2+^ level

To assess the cytosolic levels of intracellular Ca^2+^ in endothelial cells we used a ratiometric FRET-based Ca^2+^ indicator Twitch-2B, driven by the CMV promoter. Primary brain ECs were isolated and seeded in a 12-well plate on 12mm glass coverslips and maintained in an EC specific medium (Lonza, CC-3202) with or without 100μM of ATP at 37 °C and 5% CO_2_ atmosphere, replacing the growth medium every 48 hours. Transfection with CMV-Twitch-2B was performed when cell cultures were at ∼80% confluency. On the day of transfection, Lipofectamine/DNA complexes were prepared in 100μl of opti-MEM (Invitrogen) for each well of 12-well plates by mixing 1.5 μg of plasmid with 3 μl of Lipofectamine LTX and 1μl of Plus transfection reagent according to manufacturer’s instructions (Invitrogen) and added dropwise to each well (100 μl/well). Cells were incubated at 37 °C in a CO_2_ incubator for 24 h, followed by fixation in 4% PFA. The images were acquired using the CBIA Core Facility confocal microscope (LSM880 AxioObserver Z1; Zeiss) at the University of Ottawa with a 20x air objective (Plan-Apochromat 20x/NA 0.8; Zeiss). Images were acquired using a 458 nm laser to excite CFP, and the fluorescence signals were collected at the CFP and YFP emission wavelengths (461-519 and 522-578 nm bandwidth, respectively). The fluorescence signals from the regions of interest (ROIs) were calculated as a ratio of YFP to CFP emissions.

#### Analysis of Ca^2+^ transients

To assess spontaneous Ca^2+^ activity in primary mouse cerebral cortex endothelial cells we used Cal-590™ AM, a membrane-permeable organic dye that allows detection of intracellular Ca^2+^ fluctuation ^54^. ECs were cultured in an EC specific medium (Lonza, CC-3202) with or without 100μM of ATP at 37 °C and 5% CO_2_. EC medium was replaced every second day up until 80% confluence was reached (∼6 days). On the day of experiment once ECs were 80% confluent, cells were treated with 2 μM of Cal-590™ AM (AAT Bioquest, CA, USA) and were further incubated for about 1 hour in a humidified incubator at 37 °C and 5% CO_2_. After incubation, the culture dish was transferred on the stage of fluorescent microscope (DeltaVision Elite Olympus XI-71) of the CBIA Core Facility at the University of Ottawa. The microscope was equipped with an incubation chamber allowing to maintain the temperature at 37°C and CO_2_ at 5%. Images were acquired every 10 s for 5 min using a 40x oil objective, (UPLFLN 40x/NA 1.3; Olympus) and the recorded videos were analyzed using a custom-written script ^55^ in MATLAB (MathWorks). Briefly, regions of interests (ROI)s were traced around the entire cell and the fluorescence intensity in each ROI and for each time point was extracted. Ca^2+^ activity was calculated as relative changes in the percentage of ΔF/F = (F-F_back_)/ F_back_, where F is the Cal-590™ AM intensity in the ROI and F_back_ is the background signal taken in a “cell-free” region of the imaging field. Analysis of spontaneous Ca^2+^ activity was performed using a multiple threshold algorithm^56,57^. Briefly, mean standard deviation (SD) of Ca^2+^ trace for each cell was first calculated, and all peaks with an amplitude greater than 1.5 times the SD were measured. These peaks were then removed from the Ca^2+^ traces, and the mean SD of Ca^2+^ trace was re-calculated to depict events greater than 1.5 times the new SD. This procedure allowed us to depict all Ca^2+^ events regardless of their amplitudes. Ca^2+^ frequency was calculated as the number of peaks over video duration. The statistical analysis was performed in a total number of n videos collected from n wells for n cells in total in Cal-590 treated cells for 4 distinct conditions.

### Transmission electron microscopy and immunogold labeling for electron microscopy

P14 *16p11.2^df/+^* and WT mice were anesthetized using cocktail of xylazine (10mg/kg, IP) and ketamine (100 mg/kg IP). The brain was extracted and initially fixed by immersion in fixative solution (0.5% glutaraldehyde, 4% paraformaldehyde, and 0.1M sodium-cacodylate) for 1 h at RT. Tissue was then transferred to fresh fixative solution and incubated for 5 hrs at RT and then rinsed overnight with 0.1M sodium-cacodylate at 4°C. Coronal vibratome free-floating sections of 50 μm were collected and immersed in 0.1% sodium borohydride/PBS for 30 min at room temperature. Samples were thoroughly rinsed in PBS and blocked with 10% goat serum/0.5% gelatin/PBST (0.01% Triton X-100), for 2 h at room temperature. Sections were then incubated with rabbit anti-P2Y2 (1:200, Abcam: AB272891) diluted in blocking solution (without Triton X-100) overnight at room temperature. After thoroughly rinsing sections in PBS, sections were then incubated with gold-labeled goat anti-rabbit IgG (1:50, Nanoprobes: 2005-1ML), diluted in blocking solution (without Triton X-100), overnight at room temperature. Sections were washed with PBS followed by 3% sodium acetate rinse (3 × 5 min). Sections were subjected to silver-enhancement, following manufacturer’s instructions (Nanoprobes: 2012-45ML). Sections were then thoroughly rinsed in 3% sodium acetate and extensively washed in 0.1M PB pH 7.4 prior to post-fixation. Following staining, sections were post-fixed in a solution containing 1% osmium tetroxide and 1.5% potassium ferrocyanide, dehydrated by increasing ethanol concentrations followed by propylene oxide. Afterward, post-fixed sections were embedded in Durcupan ACM Epoxy resin (Electron Microscopy Sciences: 14040). Ultrathin sections (80 nm) were then cut from the block surface, using a Leica EM UC6 ultramicrotome, and collected on copper grids. Samples were examined under JEM-1400Flash transmission electron microscope operating at 80kV and equipped with a 16MP digital camera (GATAN One View).

#### Ex vivo parenchymal arteriole vascular reactivity

Following euthanasia and decapitation, mouse brains were removed and placed into cold (4°C) MOPS-buffered saline (composition: 135 mM NaCl, 5 mM KCl, 1 mM KH_2_PO_4_, 1 mM MgSO_4_, 2.5 mM CaCl_2_, 5 mM glucose, 3 mM MOPS, 0.02 mM EDTA, 2 mM pyruvate, 10 mg/mL bovine serum albumin, pH 7.3). Cortical parenchymal arterioles (PAs) originating from the middle cerebral artery (MCA) were dissected free of surrounding tissue. In an organ chamber (University of Colorado IDEA Core), PAs were then cannulated on borosilicate glass micropipettes with one end occluded, tied at both ends, and pressurized at 40mmHg using an arteriography system (Living Systems Instrumentation, Inc., St. Albans, VT, USA). PAs were perfused (4 mL/min) with prewarmed (36.5°C ± 1°C) and gassed (5% CO2, 20 % O2, 75 % N2) artificial cerebrospinal fluid (aCSF; 125 mM NaCl, 3 mM KCl, 26 mM NaHCO_3_, 1.25 mM NaH_2_PO_4_, 1 mM MgCl_2_, 4 mM glucose, 2 mM CaCl_2_, pH 7.3 at room temperature with gas aeration) for at least 30 min to allow development of myogenic tone. PA lumen diameter was continuously measured using a CCD camera and edge-detection software (IonOptix, Westwood, MA, USA). Passive PA lumen diameter was obtained in aCSF free of Ca^2+^ (0 mM [Ca^2+^]_o_ with 5 mM EGTA). PA tone was calculated with the following equation: [(passive diameter – active diameter)/(passive diameter)] × 100. Only viable PAs, defined as those that developed pressure-induced myogenic tone greater than 15% at 40 mmHg, were used in the experiments. Diameter changes of PAs, in response to bath perfusion of drug, were calculated as a percent of the maximum dilation using the following equation [(drug-induced diameter – active diameter)/(passive diameter – active diameter)]. The percent of maximum dilation in response to 10 µM bath perfusion of adenosine was calculated before and after 30-minute bath perfusion of 100 µM Diquafosol tetrasodium (DQS) in aCSF.

#### Behavioral assays

Before behavioral testing, all animals were left to habituate to an inverted light cycle housing room for 7 days. Mice were then handled and injected intraperitoneally (i.p.) with 5mg/ml of Diquafosol tetrasodium (DQS) or 1X sterile PBS as control at a volume of 1% of body weight once a day for 1 week prior to testing. Once testing was started and until testing completion, mice were injected once a day. Behavioral tests were completed at the University of Ottawa’s Behavior Core Facility between 9:00 and 17:00 under dim red light. On the testing day, before the task, animals were habituated to the testing room for 60 minutes. Behavior tests were performed with all mice in the following order: marble burying test 1 (1 day), novel-object recognition (2 days), marble burying test 2 (1 day). All tests were directly inspired from our previously published study ^6^.

##### Marble burying test

This test was performed as described elsewhere ^6^. It consisted of a 30-minute trial per mouse. For each trial, a standard polycarbonate rat cage (26 x 48 x 20cm) filled with 5-cm-thick SANI-chip bedding was used on which 20 marbles were evenly distributed (5 rows of 4 marbles). Each mouse was placed at the bottom-left corner of the cage. During the test, the cage was covered by a transparent Plexiglas. Once the trial was completed, the number of marbles that were either fully buried or at least two-thirds covered by bedding were counted as buried.

##### Novel-object recognition test

A 2-day novel-object recognition test was performed as described elsewhere ^6^. On day one, each animal was habituated to an empty open-field arena (45×45×45 cm) for 30 minutes and returned to their home cage. On the experimental day, mice were habituated for a second time to the open field for 10 minutes. Following habituation, each mouse was removed from the open field and placed in a clean holding cage for 2 minutes. Two identical objects (red cup or white funnel) were placed in the arena and the mouse returned to the arena for a 10-minute familiarization period. The mouse was removed from the arena and placed in a clean holding cage for 1hour. After the one hour, the object-recognition test consisted of one cleaned familiar object and one cleaned novel object (red cup or white funnel switched). The mouse was returned to the arena for a 5-minute recognition period. All interactions with the objects were recorded using Ethovision 17.5 XT software (Noldus). Object recognition was scored as the time during which the nose of the animal was located within 2 cm of the object. A discrimination index was calculated as follows: [time spent interacting with novel object/(time spent interacting with novel object + time spent interacting with familiar object)].

#### Statistical analyses

No statistical methods were used to predetermine sample sizes. Sample sizes were similar to those reported in previous publications ^6,7^. All sample numbers in this study are in line with well-accepted standards from the literature for each method. All data presented in this work were obtained from experimental replicates (e.g. multiple animal cohorts from different litters, at least three experimental repeats for each assay). All attempts of replication were successful. All data analyses were conducted blinded to the genotype and experimental condition. Groups were reassembled following completion of the data analysis according to genotype and experimental condition. Randomization of individual samples was performed by numbering. Statistical tests were performed using GraphPad Prism 10.0 software and MetaboAnalyst 5.0 (for metabolomics). A Mann-Whitney *U*-test or t test (appropriate for small sample sizes) was used for two-group comparisons between WT and 16p11.2-deficient ECs isolated from mice for metrics including mitochondrial density and fragmentation, mitochondrial function, Western blot quantifications and metabolite abundance. A two-way ANOVA (e.g. ‘genotype x condition’) and Sidak’s multicomparisons *post-hoc* test was used for *in vitro* network formation assay. A one-way ANOVA and Tukey’s multicomparison *post-hoc* test was used for steady state Ca^2+^ readouts and Ca^2+^ transients, behavioral assays as well as *ex vivo* vascular reactivity assessment. P<0.05 was considered significant. Statistical details of each experiment can be found in figure legends.

## SUPPLEMENTAL INFORMATION

**Figure S1.**
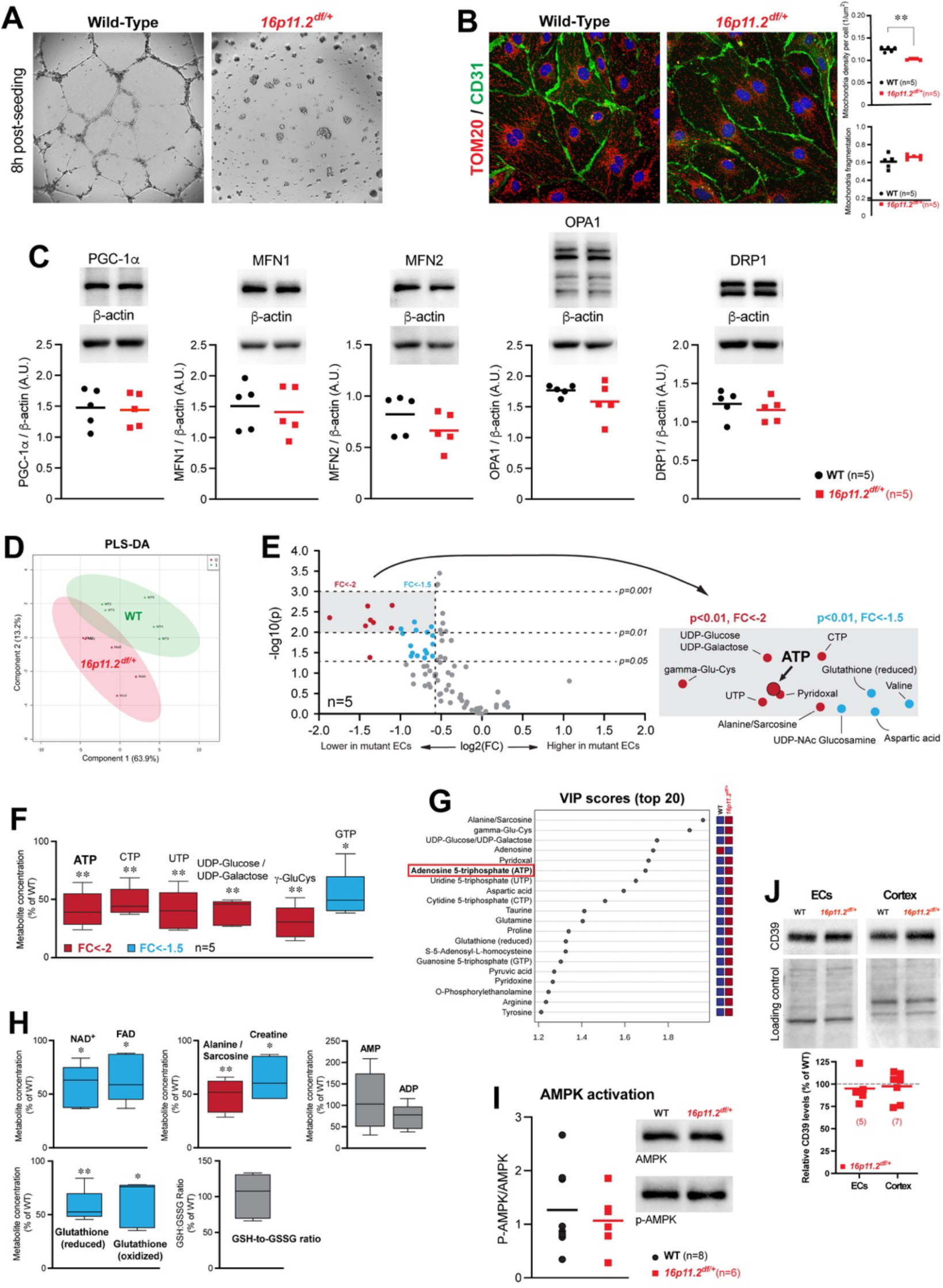
No change in mitochondrial dynamics and biogenesis in 16p11.2-deficient brain endothelial cells and additional altered metabolites, related to Figure 1. (A) Representative images of primary brain wild-type (WT) and 16p11.2-deficient ECs angiogenic ability. (B) Representative images (left) and quantification (right) of mitochondria density (top right) and mitochondria fragmentation (bottom right) in WT (n=5 mice) and 16p11.2-deficient (n=5 mice) ECs following immunostaining for mitochondrial marker TOM20 (red), endothelial marker CD31 (green) and nuclei marker DAPI (blue). \*\**p* <0.01, two-tailed Mann-Whitney test. (C) Western blot quantification of relative levels of proteins associated with mitochondrial biogenesis (PGC-1a) as well as mitochondrial fusion and fission (OPA-1, DRP1, and MFN1 and −2) in primary mouse brain endothelial cells isolated from male *16p11.2^df/+^* and WT littermates (n=5 animals). (D) Partial least-square discriminant analysis (PLS-DA) of metabolomics data from 16p11.2-deficient (n=5) and WT (n=5) ECs. Each data point represents pooled ECs isolated from three mice. Supervised PLS-DA was obtained with 2 components. The explained variances are shown in parentheses. (E) Volcano plot showing most significantly altered metabolites (p<0.01) with a fold change (FC)<-2 (red) and FC<-1.5 (blue). Plot summarizes both fold-change and t-test criteria. Scatter-plot of the negative log10-transformed p-values from the t-test plotted against the log2 fold change is shown. (F) Energy-related metabolites detected as most significantly altered by univariate statistical analysis. Metabolites displayed consist of a fold-change FC<-2 (red) or FC<-1.5 (blue). (G) Variable importance in projection (VIP) score showing the top 20 (score > 1) most important metabolites contributing to the separation of metabolic profiles identified by PLS-DA. (H) Additional metabolites found at a reduced abundance in 16p11.2-deficient ECs. Metabolites displayed consist of a fold-change (FC)<-2 (red), FC<-1.5 (blue) or unchanged (gray). (I) Western blot quantification of relative proteins associated with AMP-activated protein kinase (AMPK). No difference in AMPK activation was identified between WT (n=8) and 16p11.2-deficient (n=6) ECs. (J) Western blot quantification of relative CD39 protein levels in primary mouse brain ECs (n=5) and cerebral cortex (n=7) from 16p11.2-deficient and WT littermates. Dotted line represents 100% of WT values. (A-I) All data shown are from ECs isolated from male mice. (B, C, I and J) Data are mean with individual values. (D-H) n=5 samples, ECs pooled from 3 mice per sample. (F and H) Whisker boxes (min to max, center line indicating median) and represented as percentage of Wild-Type (WT). (F-H) \**p*< 0.05, \*\**p*<0.01, two-tailed paired *t* test.

**Figure S2.**
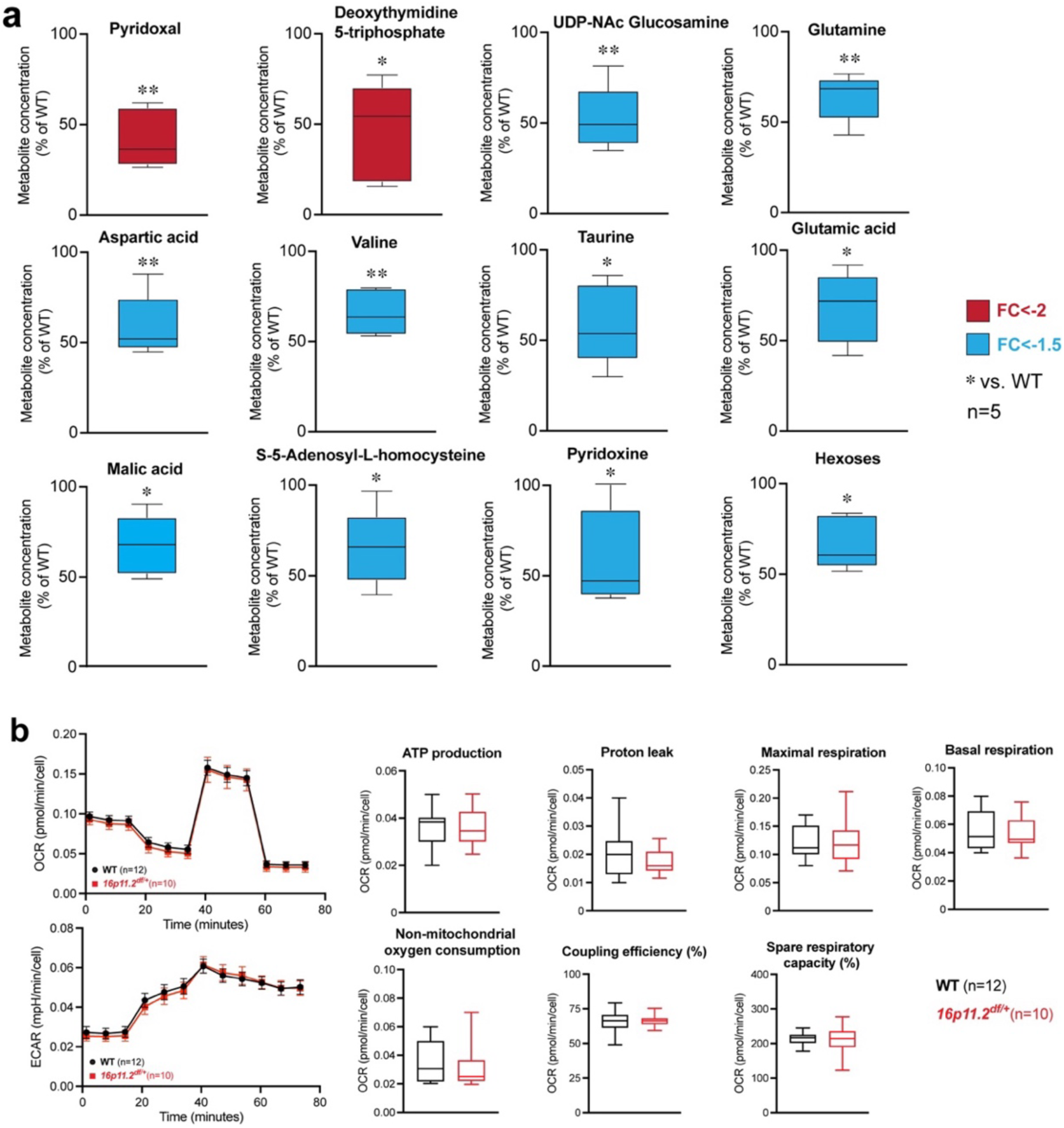
Normal mitochondrial function in 16p11.2-deficient brain endothelial cells, related to Figure 1. (A) Additional energy-related metabolites detected by univariate statistical analysis. Metabolites displayed consist of a fold-change (FC)<-2 (red), FC<-1.5 (blue) or unchanged (gray) (n=5 samples, ECs pooled from 3 mice per sample). \**p*< 0.05, \*\**p*<0.01, a two-tailed paired *t* test. (B) Quantification of oxygen consumption rate (OCR, top left) and extracellular acidification rate (ECAR, bottom left) of WT (n=12 mice) and 16p11.2-deficient (n=10 mice) ECs. Data are mean ± s.e.m. Right: quantification of additional mitochondrial functional parameters. (A and B) All data shown are from ECs isolated from male mice. Whisker boxes (min to max, center line indicating median), or traces.

**Figure S3.**
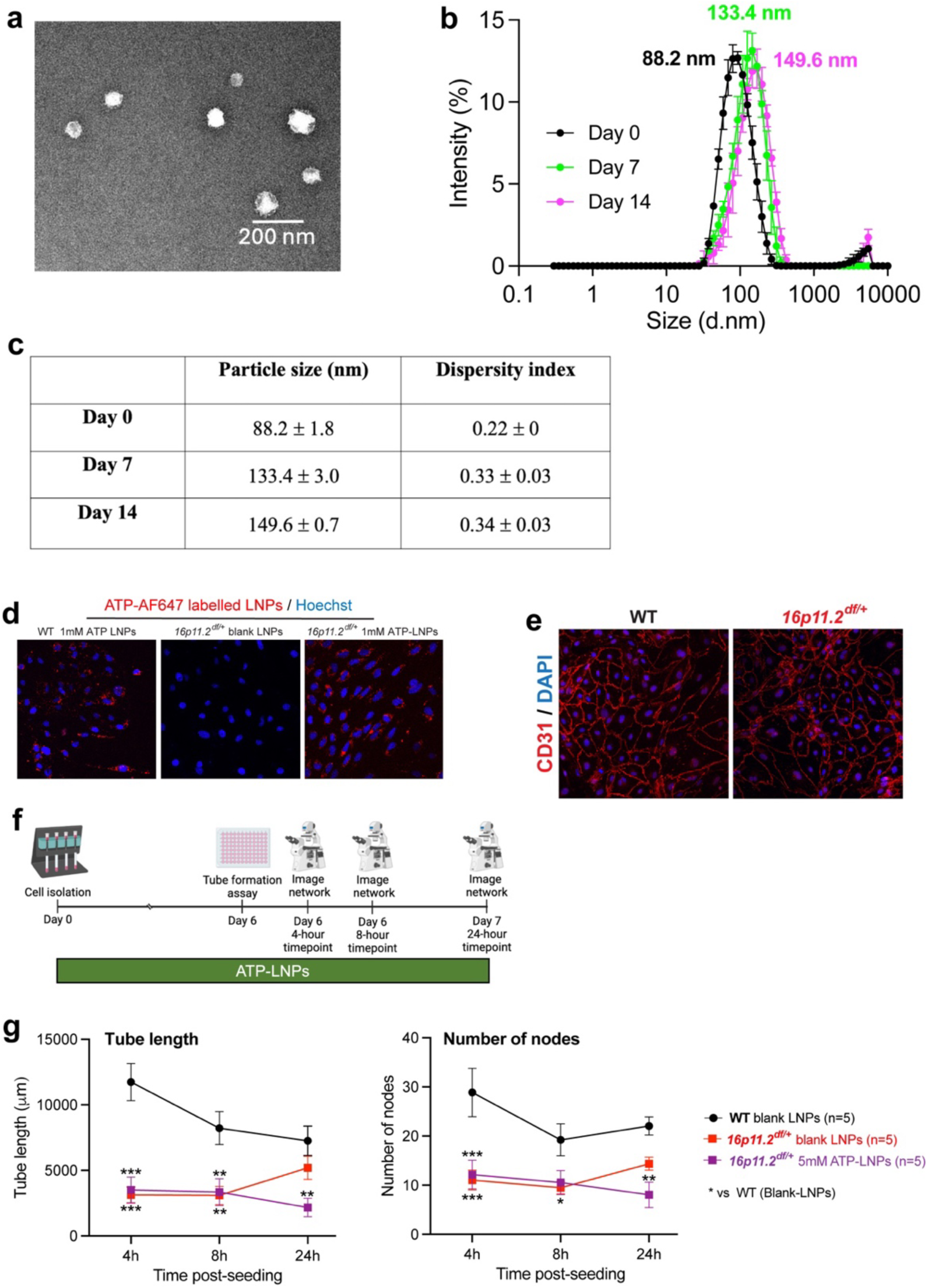
*In vitro* network formation assay of 16p11.2-deficient ECs with treatment of 5mM ATP-LNPs and confirmation of colloidal stability of ATP-LNPs, related to Figure 1. (A) Transmission electron microscopy image of ATP-LNPs. Negative-stain electron micrograph of ATP-LNPs was acquired using a Thermofisher TF20 electron microscope. Scale bar= 200 nm. (B) Intensity size distribution plots demonstrating the colloidal stability of ATP-LNPs upon storage at 4 °C for 14 days. (C) Particle size and dispersity index of LNPs using dynamic light scattering. (D) Representative images confirming intracellular delivery of ATP (red) via lipid nanoparticles (LNPs). (E) Confirmation of cell identity using immunocytochemistry with endothelial marker, CD31, (red) and nuclei marker, DAPI, (blue). (F) Experimental timeline for treatment of 16p11.2-deficient ECs with ATP-LNPs following cell isolation and during a 24-hour *in vitro* network formation assay. (G) Quantifications of network densities (total endothelial tube length) and network nodes (total number of branching hubs) following treatment of 16p11.2-deficient ECs with 5mM ATP-LNPs. Data shown are from ECs isolated from male mice and are shown as the mean ± s.e.m. (n=5 animals per group). \**p*<0.05, \*\**p*<0.01, \*\*\**p*<0.001, two-way repeated measure ANOVA and Sidak’s multicomparison *post hoc* test. (A-G) ATP-LNPs were prepared as described in Table 1. ATP-LNPs were prepared and diluted to a final ATP concentration of 40 mM using 1x PBS (pH 7.4). (B and C) Particle size and dispersity indices were measured using a Malvern Zetasizer Pro. Data are represented as mean ± SD of at least n=3 measurements.

**Figure S4.**
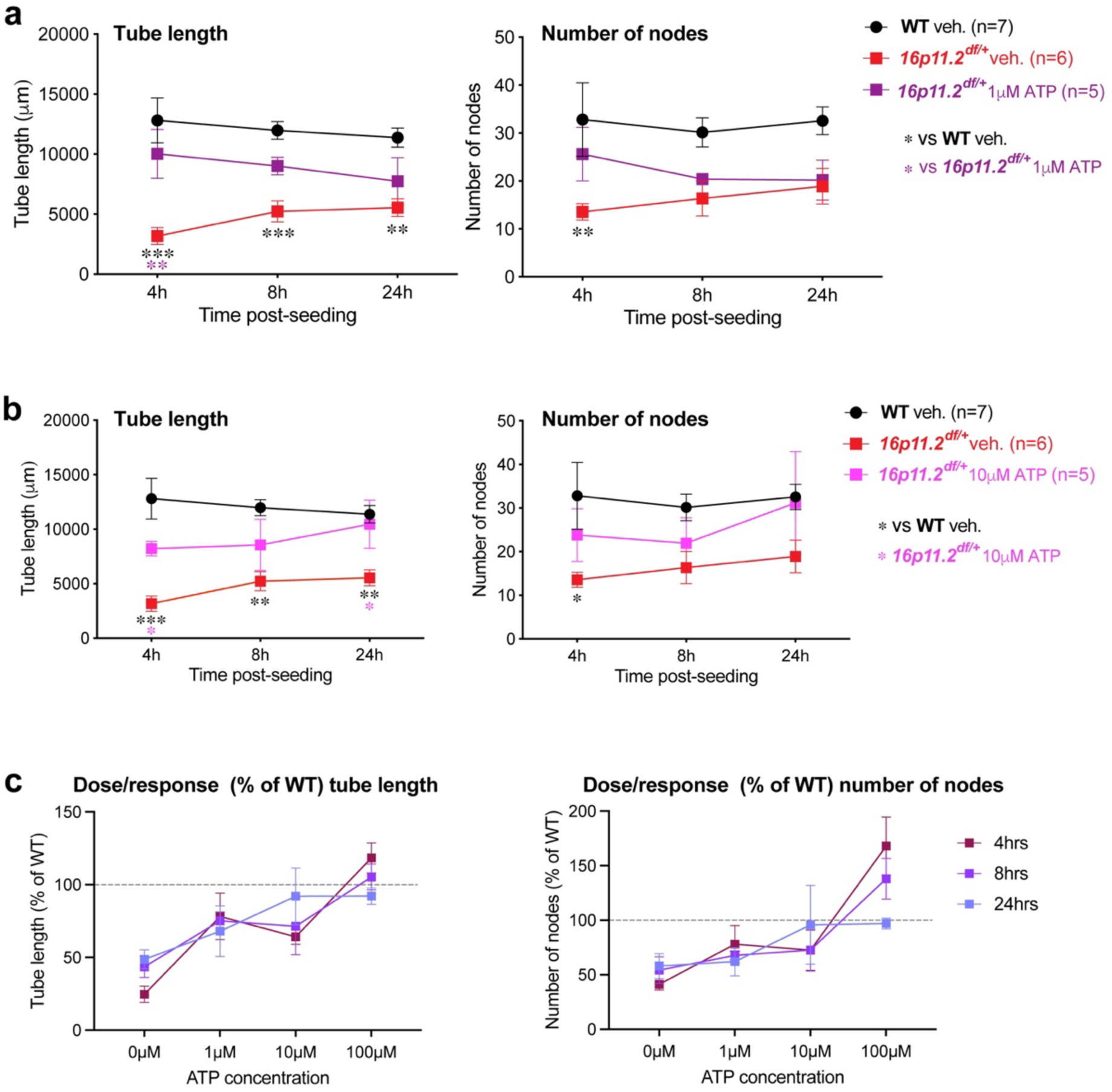
Additional treatment conditions of extracellular ATP in 16p11.2-deficient brain ECs, related to Figure 3. (A-C) Network densities (total endothelial tube length) and network nodes (total number of branching hubs) following treatment of 1µM (A) and 10µM (B) extracellular ATP (C). Dose response curves for network branching (left) and number of nodes (right) of 16p11.2-deficient ECs following extracellular ATP treatment (1µM, 10µM or 100µM) measured using an *in vitro* tube formation assay imaged at 4hrs, 8hrs and 24hrs post-seeding. Data are shown as the mean ± s.e.m. (n=5-7 animals per group). (C) Dotted line represents 100 % of WT values. (B-C) veh.= vehicle. \**p*<0.05, \*\**p*<0.01, \*\*\**p*<0.001, two-way repeated measure ANOVA and Sidak’s multicomparison *post hoc* test.

**Figure S5.**
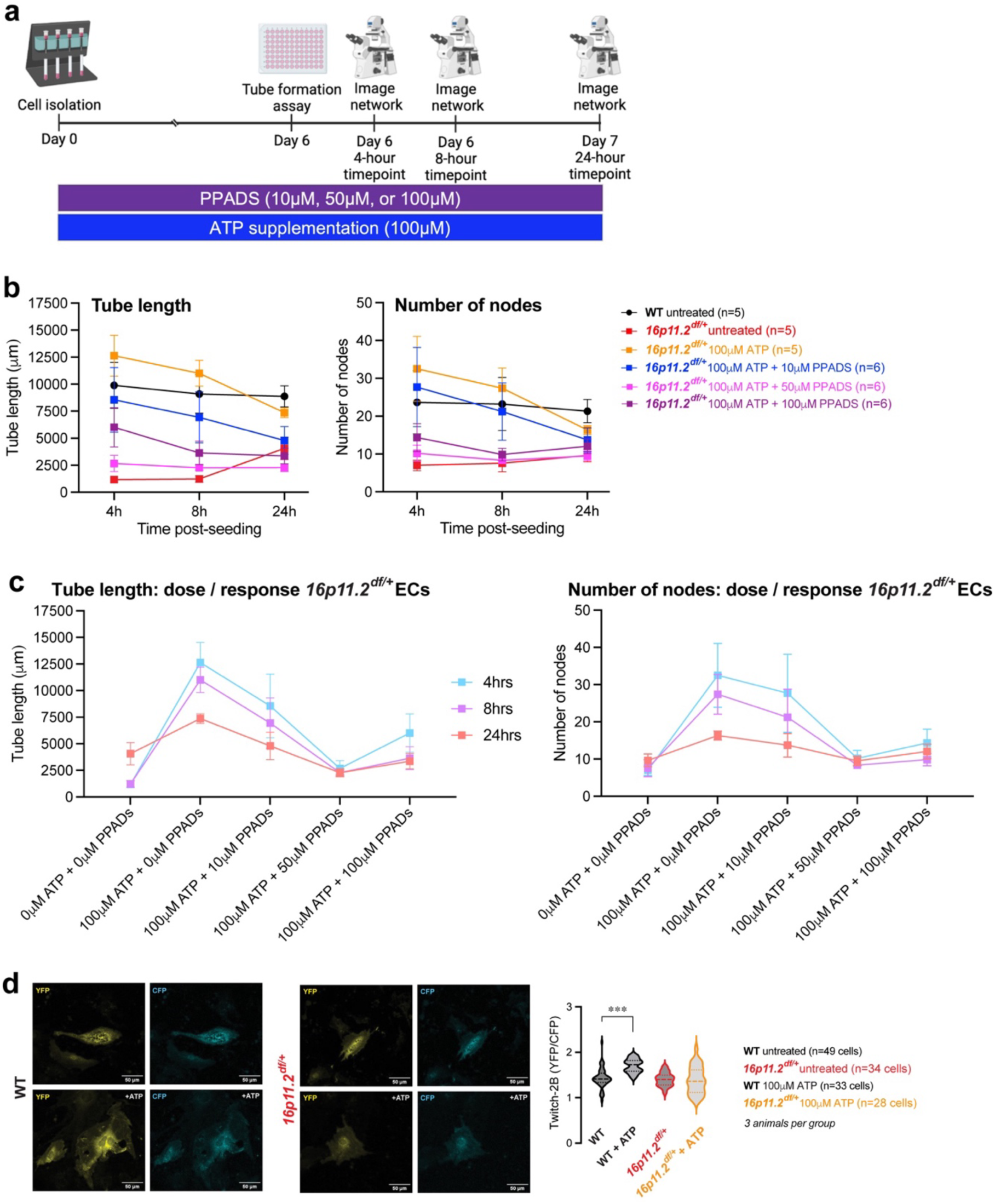
Additional treatment conditions of non-selective antagonist (PPADs) and representative dose response curves, related to Figure 3. (A) Experimental timeline for treatment of 16p11.2-deficient ECs with extracellular ATP and PPADs following cell isolation and during a 24-hour *in vitro* network formation assay. (B) Quantifications of network tube length and network nodes (total number of branching hubs) following co-administration of extracellular ATP (100µM) and PPADs (10µM, 50µM or 100µM). (C) Dose response curves of tube length (left) and number of nodes (right) of 16p11.2-deficient ECs following co-administration of extracellular ATP and PPADs measured using an *in vitro* tube formation assay imaged at 4hrs, 8hrs and 24hrs post-seeding. (D) ECs isolated from 16p11.2-deficient and WT mice were transfected with Twitch-2B, a FRET-based ratiometric Ca^2+^ indicator, to measure steady-state intracellular Ca^2+^ level. Left: representative images of yellow fluorescent protein (YFP) and cyan fluorescent protein (CFP) for both 16p11.2-deficient and WT ECs, with and without ATP are shown. Right: quantification of YFP/CFP ratio in 16p11.2-deficient and WT ECs in the absence or presence of 100 µM of extracellular ATP. Note, the lack of effect of ATP supplementation on steady-state intracellular Ca^2+^ level in 16p11.2-deficient ECs. Violin plots, center line indicating median, n=3-4 animals per group. \*\*\**p*<0.001, one-way ANOVA and Tukey multicomparison *post hoc* test. (B and C) Data are mean ± s.e.m.

**Figure S6.**
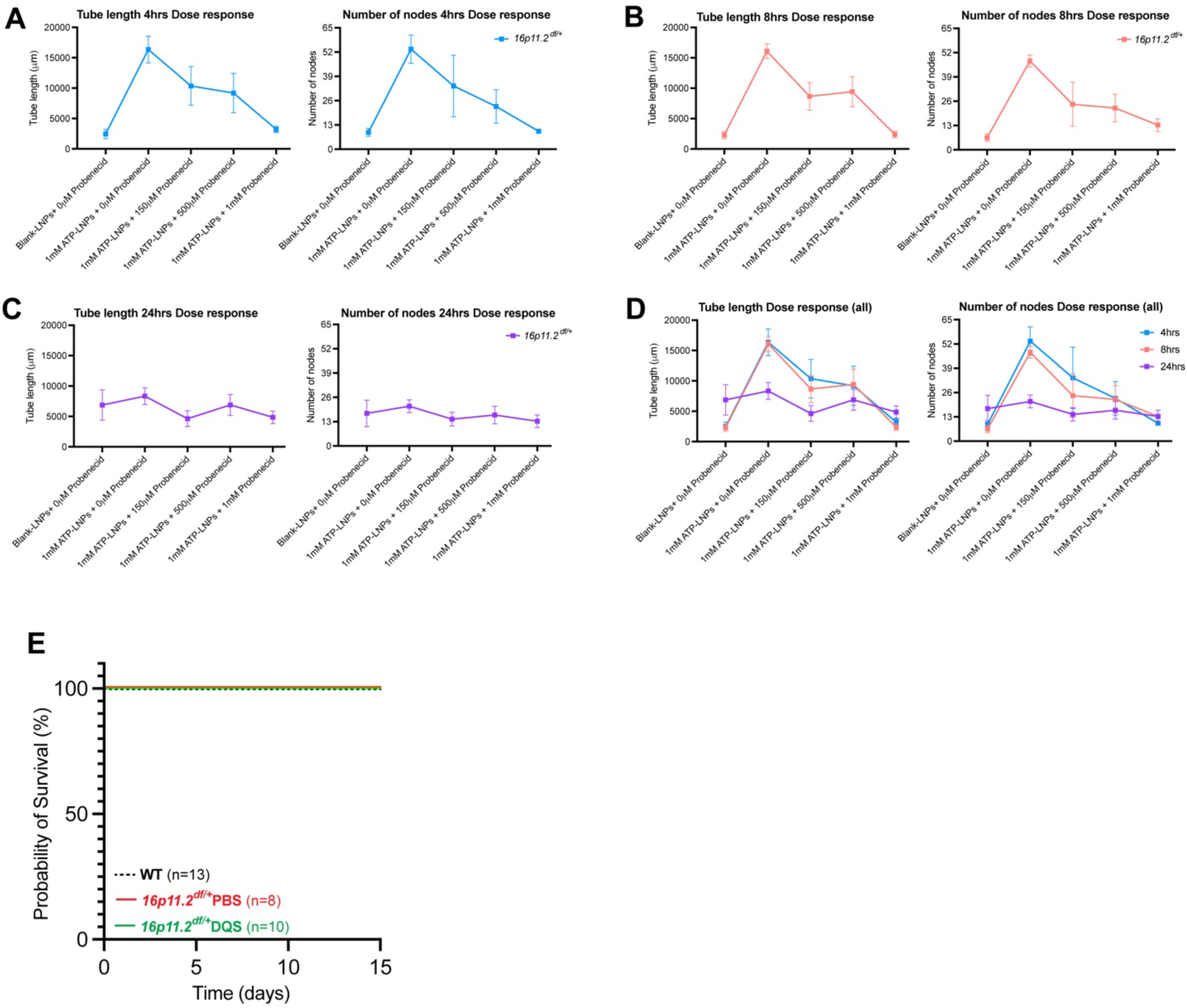
Dose response curves of 16p11.2-deficient ECs following administration of probenecid and ATP-LNPs, related to Figures 2 and 4. (A-C) Dose response curves of tube length (left) and number of nodes (right) of 16p11.2-deficient brain ECs during a 24-hour tube formation assay imaged at 4hrs (A), 8hrs (B), and 24hrs (C) post-seeding with administration of 1mM of ATP-LNPs and 150μM, 500μM or 1mM of probenecid. (D) Dose response curve summary of tube length (left) and number of nodes (right) of all imaging time-points. (E) Survival curve representing the probability of survival with daily intraperitoneal injections of Diquafosol tetrasodium (DQS). (A-D) Data are mean ± s.e.m, n=5 animals per group.

## References

1 Watts, M. E., Pocock, R. & Claudianos, C. Brain Energy and Oxygen Metabolism: Emerging Role in Normal Function and Disease. Front Mol Neurosci 11, 216 (2018). 10.3389/fnmol.2018.00216

2 Steiner, P. Brain Fuel Utilization in the Developing Brain. Ann Nutr Metab 75 Suppl 1, 8–18 (2019). 10.1159/000508054

3 Baburamani, A. A., Ek, C. J., Walker, D. W. & Castillo-Melendez, M. Vulnerability of the developing brain to hypoxic-ischemic damage: contribution of the cerebral vasculature to injury and repair? Front Physiol 3, 424 (2012). 10.3389/fphys.2012.00424

4 Vogenstahl, J., Parrilla, M., Acker-Palmer, A. & Segarra, M. Vascular Regulation of Developmental Neurogenesis. Front Cell Dev Biol 10, 890852 (2022). 10.3389/fcell.2022.890852

5 Ouellette, J. & Lacoste, B. From Neurodevelopmental to Neurodegenerative Disorders: The Vascular Continuum. Front Aging Neurosci 13, 749026 (2021). 10.3389/fnagi.2021.749026

6 Ouellette, J. et al. Vascular contributions to 16p11.2 deletion autism syndrome modeled in mice. Nat Neurosci 23, 1090–1101 (2020). 10.1038/s41593-020-0663-1

7 Beland-Millar, A. et al. 16p11.2 haploinsufficiency reduces mitochondrial biogenesis in brain endothelial cells and alters brain metabolism in adult mice. Cell Rep 42, 112485 (2023). 10.1016/j.celrep.2023.112485

8 Ouellette, J., Crouch, E. E., Morel, J. L., Coelho-Santos, V. & Lacoste, B. A Vascular-Centric Approach to Autism Spectrum Disorders. Neurosci Insights 19, 26331055241235921 (2024). 10.1177/26331055241235921

9 Vijayakumar, N. T. & Judy, M. V. Autism spectrum disorders: Integration of the genome, transcriptome and the environment. J Neurol Sci 364, 167–176 (2016). 10.1016/j.jns.2016.03.026

10 Park, H. R. et al. A Short Review on the Current Understanding of Autism Spectrum Disorders. Exp Neurobiol 25, 1–13 (2016). 10.5607/en.2016.25.1.1

11 Bosl, W. J., Tager-Flusberg, H. & Nelson, C. A. EEG Analytics for Early Detection of Autism Spectrum Disorder: A data-driven approach. Sci Rep 8, 6828 (2018). 10.1038/s41598-018-24318-x

12 Takarae, Y. & Sweeney, J. Neural Hyperexcitability in Autism Spectrum Disorders. Brain Sci 7 (2017). 10.3390/brainsci7100129

13 Courchesne, E. et al. Neuron Number and Size in Prefrontal Cortex of Children With Autism. JAMA 306, 2001–2010 (2011).

14 Kaushik, G. & Zarbalis, K. S. Prenatal Neurogenesis in Autism Spectrum Disorders. Frontiers in Chemistry 4 (2016).

15 Donovan, A. P. & Basson, M. A. The neuroanatomy of autism - a developmental perspective. J Anat 230, 4–15 (2017). 10.1111/joa.12542

16 Lacoste, B. et al. Sensory-Related Neural Activity Regulates the Structure of Vascular Networks in the Cerebral Cortex. Neuron 83, 1117–1130 (2014). 10.1016/j.neuron.2014.07.034

17 Owen, J. B. a. B., A. Measurement of Oxidized/Reduced Glutathione Ratio. Protein Misfolding and Cellular Stress in Disease and Aging, 269–277 (2010).

18 Bjorklund, G. et al. The role of glutathione redox imbalance in autism spectrum disorder: A review. Free Radic Biol Med 160, 149–162 (2020). 10.1016/j.freeradbiomed.2020.07.017

19 Muraoka, M. et al. Reactivity of gamma-glutamyl-cysteine with intracellular and extracellular glutathione metabolic enzymes. FEBS Lett 596, 180–188 (2022). 10.1002/1873-3468.14261

20 Grahame Hardie, D. AMP-activated protein kinase: a key regulator of energy balance with many roles in human disease. J Intern Med 276, 543–559 (2014). 10.1111/joim.12268

21 Wilson, C., Lee, M. D., Buckley, C., Zhang, X. & McCarron, J. G. Mitochondrial ATP Production is Required for Endothelial Cell Control of Vascular Tone. Function (Oxf) 4, zqac063 (2023). 10.1093/function/zqac063

22 Khare, P., Conway, J. F. & D, S. M. Lipidoid nanoparticles increase ATP uptake into hypoxic brain endothelial cells. Eur J Pharm Biopharm 180, 238–250 (2022). 10.1016/j.ejpb.2022.10.011

23 Chen, S. et al. Influence of particle size on the in vivo potency of lipid nanoparticle formulations of siRNA. J Control Release 235, 236–244 (2016). 10.1016/j.jconrel.2016.05.059

24 Cullis, P. R. & Hope, M. J. Lipid Nanoparticle Systems for Enabling Gene Therapies. Mol Ther 25, 1467–1475 (2017). 10.1016/j.ymthe.2017.03.013

25 Lohman, A. W., Billaud, M. & Isakson, B. E. Mechanisms of ATP release and signalling in the blood vessel wall. Cardiovasc Res 95, 269–280 (2012). 10.1093/cvr/cvs187

26 Wang, Y. et al. Purinergic signaling: A gatekeeper of blood-brain barrier permeation. Front Pharmacol 14, 1112758 (2023). 10.3389/fphar.2023.1112758

27 Thakore, P. et al. Brain endothelial cell TRPA1 channels initiate neurovascular coupling. Elife 10 (2021). 10.7554/eLife.63040

28 Ray, A., Zoidl, G., Weickert, S., Wahle, P. & Dermietzel, R. Site-specific and developmental expression of pannexin1 in the mouse nervous system. Eur J Neurosci 21, 3277–3290 (2005). 10.1111/j.1460-9568.2005.04139.x

29 Muhleder, S. et al. Purinergic P2Y(2) receptors modulate endothelial sprouting. Cell Mol Life Sci 77, 885–901 (2020). 10.1007/s00018-019-03213-2

30 Bintig, W. et al. Purine receptors and Ca(2+) signalling in the human blood-brain barrier endothelial cell line hCMEC/D3. Purinergic Signal 8, 71–80 (2012). 10.1007/s11302-011-9262-7

31 Khakh, B. S. & North, R. A. Neuromodulation by extracellular ATP and P2X receptors in the CNS. Neuron 76, 51–69 (2012). 10.1016/j.neuron.2012.09.024

32 Dai, S. et al. Purine signaling pathway dysfunction in autism spectrum disorders: Evidence from multiple omics data. Front Mol Neurosci 16, 1089871 (2023). 10.3389/fnmol.2023.1089871

33 Yamazaki, K., Yoneyama, J., Kimoto, R., Shibata, Y. & Mimura, T. Prevention of Surgery-Induced Dry Eye by Diquafosol Eyedrops after Femtosecond Laser-Assisted Cataract Surgery. J Clin Med 11 (2022). 10.3390/jcm11195757

34 Kim, S., Shin, J. & Lee, J. E. A randomised, prospective study of the effects of 3% diquafosol on ocular surface following cataract surgery. Sci Rep 11, 9124 (2021). 10.1038/s41598-021-88589-7

35 Angelakos, C. C. et al. Hyperactivity and male-specific sleep deficits in the 16p11.2 deletion mouse model of autism. Autism Res 10, 572–584 (2017). 10.1002/aur.1707

36 Portmann, T. et al. Behavioral abnormalities and circuit defects in the basal ganglia of a mouse model of 16p11.2 deletion syndrome. Cell Rep 7, 1077–1092 (2014). 10.1016/j.celrep.2014.03.036

37 Glinton, K. a. E., S. Untargeted Metabolomics for Autism Spectrum Disorders: Current Status and Future Directions. Front in Psychiatry 10 (2019). 10.3389/fpsyt.2019.00647published10.3389fpsyt.2019.00647Untargeted

38 Likhitweerawong, N. et al. Profiles of urine and blood metabolomics in autism spectrum disorders. Metab Brain Dis 36, 1641–1671 (2021). 10.1007/s11011-021-00788-3

39 Lingampelly, S. S. et al. Metabolic network analysis of pre-ASD newborns and 5-year-old children with autism spectrum disorder. Commun Biol 7, 536 (2024). 10.1038/s42003-024-06102-y

40 Khaliulin, I., Hamoudi, W. & Amal, H. The multifaceted role of mitochondria in autism spectrum disorder. Mol Psychiatry 30, 629–650 (2025). 10.1038/s41380-024-02725-z

41 Lin, R. Z. et al. Mitochondrial transfer mediates endothelial cell engraftment through mitophagy. Nature 629, 660–668 (2024). 10.1038/s41586-024-07340-0

42 Losenkova, K. et al. Endothelial cells cope with hypoxia-induced depletion of ATP via activation of cellular purine turnover and phosphotransfer networks. Biochim Biophys Acta Mol Basis Dis 1864, 1804–1815 (2018). 10.1016/j.bbadis.2018.03.001

43 Burnstock, G. & Ralevic, V. Purinergic signaling and blood vessels in health and disease. Pharmacol Rev 66, 102–192 (2014). 10.1124/pr.113.008029

44 Wang, S. et al. P2Y(2) and Gq/G(1)(1) control blood pressure by mediating endothelial mechanotransduction. J Clin Invest 125, 3077–3086 (2015). 10.1172/JCI81067

45 Ju, J. et al. Adenosine mediates the amelioration of social novelty deficits during rhythmic light treatment of 16p11.2 deletion female mice. Mol Psychiatry (2024). 10.1038/s41380-024-02596-4

46 Babiec, L., Wilkaniec, A., Gawinek, E., Hilgier, W. & Adamczyk, A. Inhibition of purinergic P2 receptors prevents synaptic and behavioral alterations in a rodent model of autism spectrum disorders. Research in Autism Spectrum Disorders 112 (2024). 10.1016/j.rasd.2024.102353

47 Lanser, A. J. et al. Disruption of the ATP/adenosine balance in CD39(-/-) mice is associated with handling-induced seizures. Immunology 152, 589–601 (2017). 10.1111/imm.12798

48 Fu, Z. et al. Microglia modulate the cerebrovascular reactivity through ectonucleotidase CD39. Nat Commun 16, 956 (2025). 10.1038/s41467-025-56093-5

49 Horev, G. et al. Dosage-dependent phenotypes in models of 16p11.2 lesions found in autism. Proc Natl Acad Sci U S A 108, 17076–17081 (2011). 10.1073/pnas.1114042108

50 Lindeberg, T. Feature detection with automatic scale selection. International journal of computer vision 30, 79–116 (1998).

51 Freitas-Andrade, M. et al. Astroglial Hmgb1 regulates postnatal astrocyte morphogenesis and cerebrovascular maturation. Nat Commun 14, 4965 (2023). 10.1038/s41467-023-40682-3

52 Whitehead, K. A. et al. Degradable lipid nanoparticles with predictable in vivo siRNA delivery activity. Nature communications 5, 4277 (2014).

53. Carpentier, G. Angiogenesis Analyzer for ImageJ, <http://image.bio.methods.free.fr/ImageJ/?Angiogenesis-Analyzer-for-ImageJ#outil_sommaire_0> (2012).

54 Tischbirek, C. H., Birkner, A. & Konnerth, A. In vivo deep two-photon imaging of neural circuits with the fluorescent Ca(2+) indicator Cal-590. J Physiol 595, 3097–3105 (2017). 10.1113/JP272790

55 Ropa, J., Cooper, S., Capitano, M. L., Van’t Hof, W. & Broxmeyer, H. E. Human Hematopoietic Stem, Progenitor, and Immune Cells Respond Ex Vivo to SARS-CoV-2 Spike Protein. Stem Cell Rev Rep 17, 253–265 (2021). 10.1007/s12015-020-10056-z

56 Gengatharan, A. et al. Adult neural stem cell activation in mice is regulated by the day/night cycle and intracellular calcium dynamics. Cell 184, 709–722 e713 (2021). 10.1016/j.cell.2020.12.026

57 Malvaut, S. et al. CaMKIIalpha Expression Defines Two Functionally Distinct Populations of Granule Cells Involved in Different Types of Odor Behavior. Curr Biol 27, 3315–3329 e3316 (2017). 10.1016/j.cub.2017.09.058

